# Functional Coupling of TRPM2 and NMDARs exacerbates excitotoxicity in ischemic brain injury

**DOI:** 10.1101/2021.07.29.454247

**Authors:** Pengyu Zong, Jianlin Feng, Zhichao Yue, Gongxiong Wu, Baonan Sun, Yanlin He, Barbara Miller, Albert S. Yu, Zhongping Su, Yasuo Mori, Jia Xie, Lixia Yue

## Abstract

Excitotoxicity caused by NMDA receptors (NMDARs) is a major cause of neuronal death in ischemic stroke. However, past efforts of directly targeting NMDARs have unfortunately failed in clinical ischemic stroke trials. Here we reveal an unexpected mechanism underlying NMDARs-mediated neurotoxicity, which leads to identification of a novel target and development of an effective therapeutic peptide for ischemic stroke. We show that NMDAR’s excitotoxicity upon ischemic insults is mediated by physical and functional coupling to TRPM2. The physical interaction of TRPM2 with NMDARs results in markedly increase in the surface expression of NMDARs, leading to enhanced NMDAR function and increased neuronal death. We identified a specific NMDAR-interacting domain on TRPM2, and developed a cell-permeable peptide to uncouple TRPM2-NMDARs. The disrupting-peptide protects neurons against ischemic injury *in vitro* and protects mice against ischemic stroke *in vivo*. These findings provide an unconventional strategy to eliminate excitotoxic neuronal death without directly targeting NMDARs.

**HIGHLIGHTS:**

- TRPM2 physically and functionally interacts with NMDARs
- Interaction of TRPM2 with NMDARs exacerbates NMDAR’s extrasynaptic excitotoxicity by increasing NMDAR’s surface expression during ischemic injury
- TRPM2 recruits PKCγ to the interacting complexes to increase NMDAR’s surface expression
- Uncoupling the interaction between TRPM2 and NMDARs with a disrupting peptide (TAT-EE_3_) protects neurons against ischemic stroke *in vitro* and *in vivo*

**GRAPHIC ABSTRACT:** 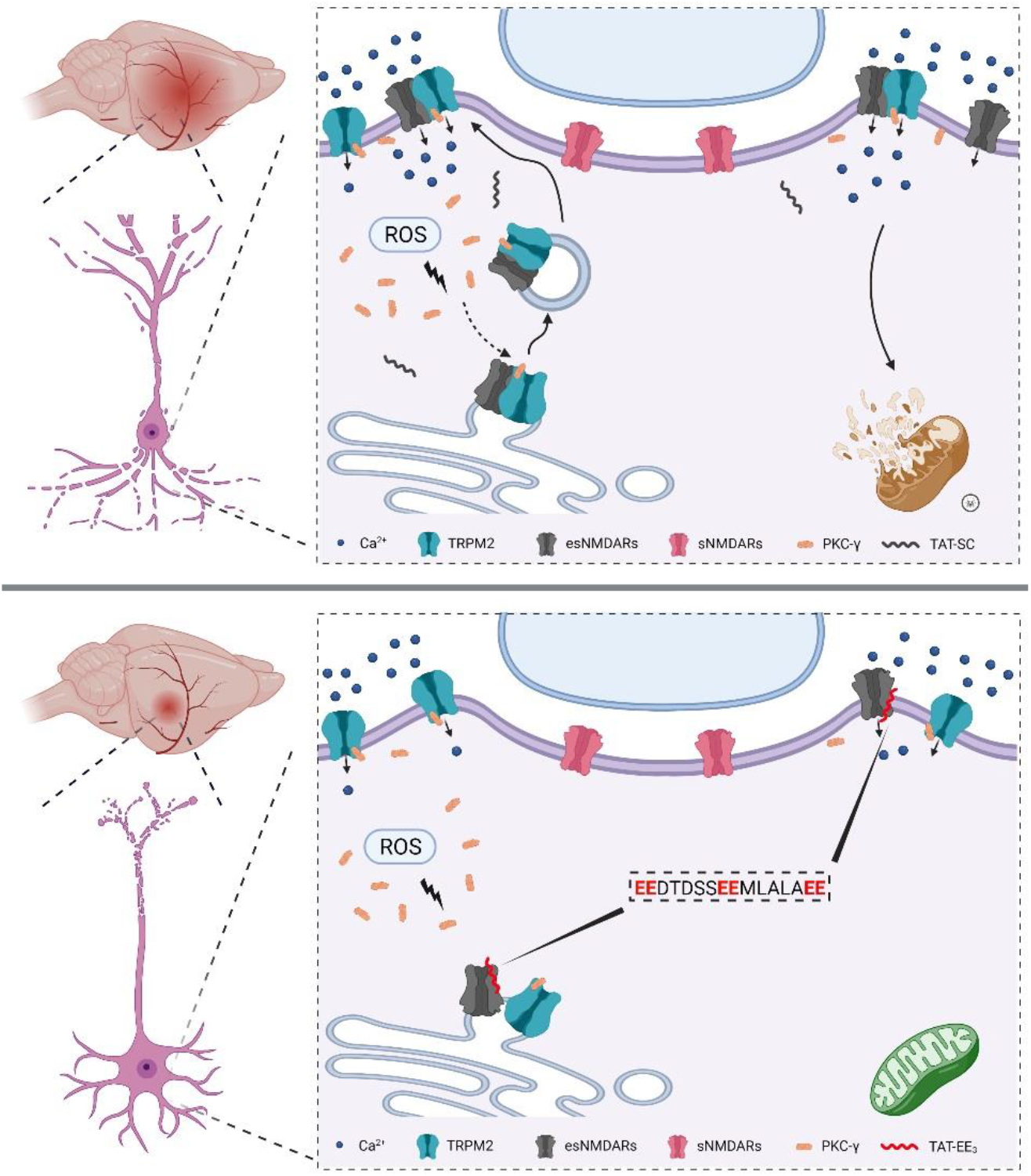

TRPM2 excerbates NMDAR’s excitotoxicity by physically and functionally interacting with NMDARs. The disrupting pipette TAT-EE_3_ protects neurons against ischemic injury *in vitro* and *in vivo*.

## INTRODUCTION

Neuronal death is a hallmark of ischemic stroke, a devastating neurological disease which remains a leading cause of disability and mortality worldwide (Virani et al., 2020). Numerous factors are involved in neuronal damage during ischemic stroke, among which Ca^2+^ overload plays a key role in neurotoxicity (Granzotto et al., 2020). Ca^2+^ overload caused by excitotoxic mechanisms through NMDA receptor (NMDAR) activation and non-excitotoxic Ca^2+^ entry mechanisms triggers a series of downstream signaling cascades, including reactive oxygen species (ROS) or reactive nitrogen species (RNS) generation, mitochondrial dysfunction, metabolic impairment, and activation of necrosis and apoptosis cascade, and ultimately leads to neuronal death (Choi, 2020). Since it was first discovered fifty years ago (Olney, 1969), excitotoxicity caused by NMDAR-mediated Ca^2+^ overload has been the center of extensive research for understanding the underlying mechanisms and for developing effective therapeutics for ischemic stroke. However, the development of stroke drugs by antagonizing NMDARs has been characterized by success in animal studies but subsequent failure in clinical trials (Sena et al., 2007).

The lack of clinical success with excitotoxic NMDAR antagonists prompted a shift of the focus of stroke neuroprotection research towards the identification of downstream intracellular signaling pathways triggered by NMDARs (Wu and Tymianski, 2018), and the investigation of subtype-dependent (Ge et al., 2020) as well as localization-dependent excitotoxic effects of NMDARs (Hardingham and Bading, 2010). Over a third of surface NMDARs are located extrasynaptically (Petit-Pedrol and Groc, 2021), which preferentially leads to neurotoxicity and cell death upon activation, whereas activation of synaptic NMDARs promotes a survival mechanism, likely through activation of differential signaling pathways triggered by intracellular Ca^2+^ (Bading, 2013; Hardingham, 2019). Moreover, the disappointing clinical trial outcome of NMDAR antagonists for stroke treatment also prompted a divergent focus on investigating non-excitotoxic Ca^2+^-permeable channels as potential therapeutic targets (Tymianski, 2011), including the Ca^2+^-permeable nonselective transient receptor potential (TRP) channels such as TRPM2 (Belrose and Jackson, 2018).

TRPM2 was discovered as an oxidative stress-activated Ca^2+^-permeable non-selective cation channel (Hara et al., 2002; Perraud et al., 2001; Sano et al., 2001), belonging to the TRP superfamily, melastatin subfamily (Clapham, 2003; Montell et al., 2002). A rise of intracellular Ca^2+^ ([Ca^2+^]_i_) and the binding of ADP ribose (ADPR) to the N-terminal binding site (Huang et al., 2019; Huang et al., 2018; Kuhn et al., 2016), and/or C-terminal NUDT9-H domain (Csanady and Torocsik, 2009; Kuhn and Luckhoff, 2004; Perraud et al., 2001; Wang et al., 2018; Yu et al., 2017), triggers conformational changes and opens TRPM2. TRPM2 is susceptible to, and regulated by, stress conditions such as acidic intra- and extracellular pH (Du et al., 2009b; Starkus et al., 2010; Yang et al., 2010), glutathione (GSH) (Belrose et al., 2012), and Zn^2+^ (Mortadza et al., 2017). TRPM2 is also sensitive to temperature (Kashio et al., 2012) and is involved in body temperature sensation (Kashio and Tominaga, 2017; Song et al., 2016; Tan and McNaughton, 2016; Vilar et al., 2020). The ADPR and Ca^2+^-gating features make TRPM2 a common molecular mechanism conferring the susceptibility to cell death induced by ROS and diverse pathological conditions.

TRPM2 is ubiquitously expressed in various cell types and most abundantly in the brain (Fonfria et al., 2006). In response to oxidative stress stimuli, TRPM2-mediated Ca^2+^ influx leads to cell death of various cell types including neurons (Belrose and Jackson, 2018; Mai et al., 2020; Takahashi et al., 2011). Contribution of TRPM2 to ischemic brain stroke has been demonstrated in ischemia-reperfusion brain damage mouse models (Alim et al., 2013; Gelderblom et al., 2014; Shimizu et al., 2013). However, the reported results have been controversial regarding whether TRPM2 in neurons or TRPM2 in immunocompetent cells plays a key role in causing ischemia-reperfusion brain damage (Alim et al., 2013; Gelderblom et al., 2014; Shimizu et al., 2013), and whether TRPM2 inhibition or knockdown (Jia et al., 2011; Shimizu et al., 2013) only protects against ischemic brain damage in male mice. Nonetheless, although the mechanisms by which TRPM2 results in deleterious effects during ischemic stroke require further investigation, TRPM2 has been implicated as a non-excitotoxic candidate target for ischemic stroke (Belrose and Jackson, 2018; Mai et al., 2020).

Given the complexity of deleterious effects caused by both excitotoxic Ca^2+^ and non-excitotoxic Ca^2+^ signaling pathways during ischemic stroke, blocking just one of these pathways may not be effective in mitigating ischemic injury. We reasoned that a convergent inhibition of the divergent excitotoxic and non-excitotoxic pathways might produce a better therapeutic outcome. Here we unveil a previously unknown mechanism by which TRPM2 mediates deleterious effects leading to neuronal death during ischemic stroke. We found that TRPM2 exacerbates NMDARs’ excitotoxicity by physically and functionally interacting with NMDARs triggered by oxidative stress. By discovering the binding domain and designing a disruptive peptide TAT-EE_3_, we demonstrated that functional uncoupling of TRPM2 and NMDARs by TAT-EE_3_ protects neurons against ischemic injury *in vitro* and *in vivo*. Our results establish that TRPM2 is a molecule converging the excitotoxic and non-excitotoxic pathways. Targeting TRPM2 represents a new therapeutic strategy to eliminate excitotoxicity caused by NMDARs in ischemic stroke.

## RESULTS

### TRPM2 deletion in neurons prevents ischemic injury and protects the brain against ischemic stroke

The oxidative stress-activated TRPM2 is expressed in various types of cells (Fonfria et al., 2006). Inhibition of TRPM2 attenuates ischemic injury, yet the mechanism by which TRPM2 leads to deleterious effects is not fully understood, as both neurons and immune cells were suggested as the primary cause of ischemic injury mediated by TRPM2 (Alim et al., 2013; Gelderblom et al., 2014). To elucidate the underlying mechanisms and determine neuronal damage mediated by TRPM2 during ischemic stroke, we established a neuron-specific *Trpm2* deletion model using nestin-cre mice crossed with TRPM2^fl/fl^ mice. Global knockout of TRPM2 (TRPM2-KO) was used as a comparison. *Trpm2* deletion was confirmed by detecting TRPM2 protein expression and functional current recording (Figure S1). Using a 120-min middle cerebral artery occlusion (MCAO) followed by reperfusion, we evaluated infarct volume 24 hrs after MCAO by TTC staining. Successful MCAO was confirmed by monitoring blood flow reduction by 85% (Figure S2). **Figure 1** shows that, similar to the protective effects produced by global *Trpm2* knockout (gM2KO) as previously reported (Alim et al., 2013) (Figure 1A-C), neuron-specific *Trpm2* deletion (Cre^+^, TRPM2^fl/fl^; nM2KO) exhibited significantly reduced infarct volume and markedly improved neurological performance in comparison with wild-type (Cre^-^, TRPM2^fl/fl^; WT) littermates (Figure 1D-F). These results establish that TRPM2 in neurons play a key role in mediating neuronal cell death. To further determine the mechanisms of TRPM2-mediated neuronal death during ischemic stroke, we evaluated neuronal cell death by TUNEL staining. At the ischemic penumbra, the numbers of TUNEL-positive neurons were significantly smaller in global TRPM2-KO and neuron-specific TRPM2-KO brain slices than those of WT littermates (Figure 1G-L), indicating that attenuation of apoptosis by *Trpm2* deletion mediates protective effects against ischemic stroke.

**Figure 1.**
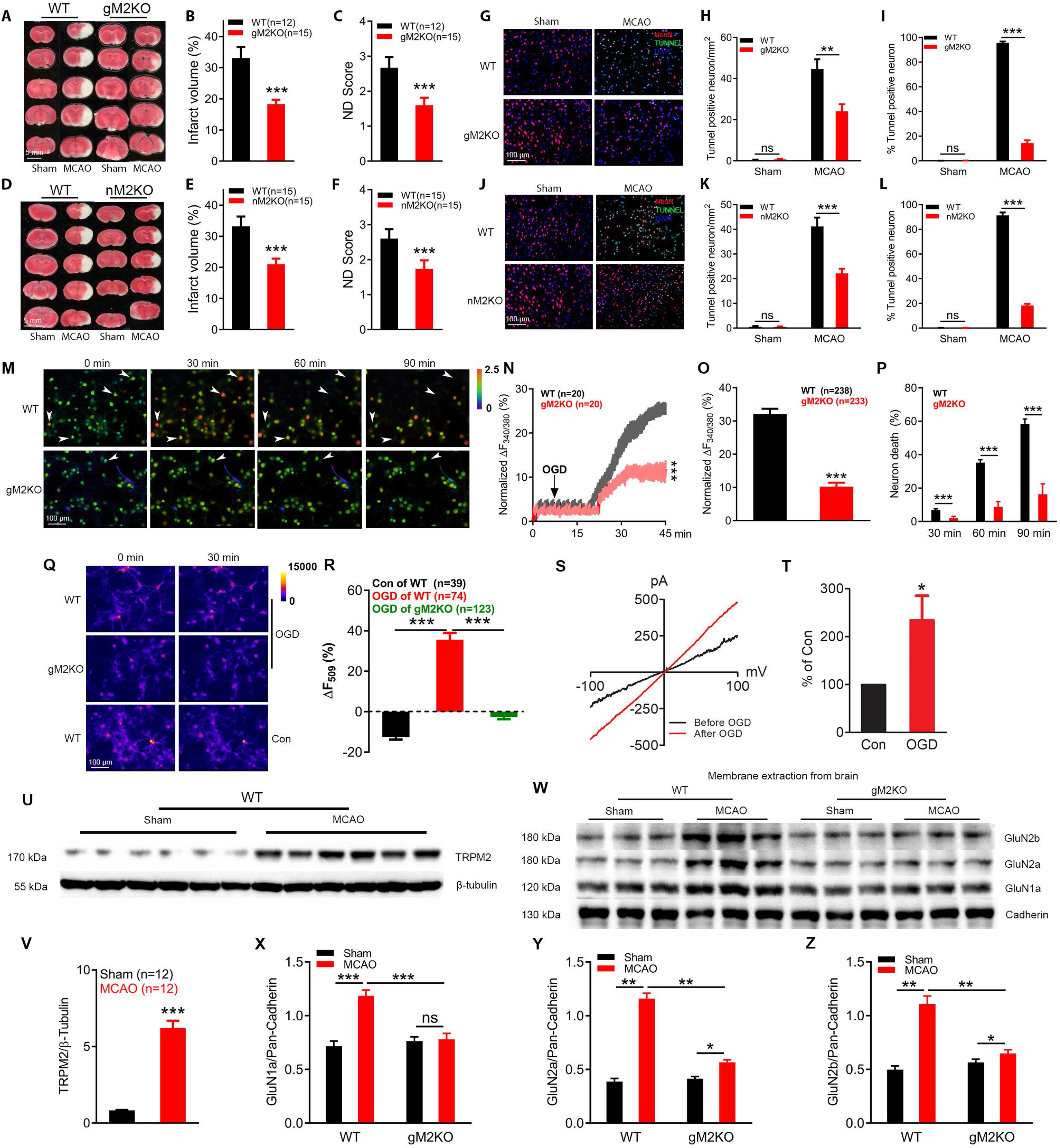
Neuron-specific *Trpm2* knockout protects the brain against ischemic damage via reducing excitotoxicity during ischemia stroke. (A-C), Global *Trpm2* deletion (gM2KO) reduces infarct volume and improves neurological deficit (ND) score in mice subjected to MCAO. In (A), representative images of TTC staining of slices at 1 mm thickness from brains of wild-type (WT) and gM2KO mice 24 hrs after sham or MCAO surgery. In (B), mean infarct volume after MCAO from WT (n=12) and gM2KO brains (n=15). In (C), average ND score 24 hrs after MCAO from 12 WT and 15 gM2KO mice (***: p < 0.001; ANOVA, Bonferroni’s test; mean ± SEM). (D-F), Neuron-specific *Trpm2* deletion by nestin Cre (Cre^+^: nM2KO for simplicity) produces similar protective effects to that of gM2KO for ischemic stroke as shown in (A-C). (D), Representative images of TTC staining of Cre^+^ *Trpm2* knockout mice (nM2KO) and Cre^-^ control littermates (WT: for simplicity) 24 hrs after sham or MCAO surgery. (E), Average infarct volume after MCAO from 15 WT and 15 nM2KO mouse brains (***, p < 0.001; ANOVA, Bonferroni’s t-test, mean ± SEM). (F), Mean ND score of the 15 nM2KO and WT mice 24 hrs after MCAO. (G-L), Neuronal death evaluated by TUNEL staining of brain sections from gM2KO versus WT mice (G-I) and nM2KO versus Cre^-^ control WT littermates. (G, J), Representative merged images of TUNEL staining of brain sections of WT and M2KO (G), and brain sections of nM2KO and WT 24 hrs after MCAO or sham surgery (Red: NeuN; Blue: DAPI; Green: TUNEL). (H, K), Quantification of TUNEL-positive neurons of WT and gM2KO (H), as well as nM2KO and WT sections from 5 mice/group (**, p < 0.01; ANOVA, Bonferroni’s t-test, mean ± SEM). (I, L), Mean percentage of TUNNEL positive neurons in all NeuN positive cells (***, p < 0.001; ANOVA, Bonferroni’s test; mean ± SEM). M-P, Evaluation of Ca^2+^ overload and neuronal death caused by OGD using ratio Ca^2+^ imaging. Cortical neurons isolated from WT and gM2KO mice were cultured for 7 to 14 days before OGD experiments. (M), Ratio Ca^2+^ imaging showing intracellular Ca^2+^ changes induced by OGD in WT and gM2KO neurons. Fluorescence ratio F_340/380_ was used to represent intracellular Ca^2+^ changes (scale bar in bottom left image = 100μm). Neurons with increasingly elevated Ca^2+^ levels, such as the ones indicated with arrows, die with time, as reflected by the disappearance of the fluorescence at the next time point. Ionomycin was used to induce the maximum Ca^2+^ influx for normalization (not shown). (N), Representative real-time changes of Ca^2+^ induced by OGD in the first 45 min. The averaged traces were from 20 neurons randomly chosen from a representative culture dish of WT and gM2KO groups (***, p < 0.001, unpaired *t*-test, mean ± SEM). (O), Quantification of OGD-induced Fura-2 fluorescence changes 30 min after OGD. A cohort of 238 neurons from three WT mice in 6 culture dishes and 233 neurons from gM2KO mice in 6 culture dishes were used for analysis (***, p < 0.001, unpaired *t*-test, mean ± SEM). (P), OGD-induced neuronal death at 30 min, 60 min, and 90 min after OGD (***, p < 0.001, unpaired *t*-test, mean ± SEM, n=238 and 233 neurons in WT and gM2KO groups). Neuronal death was monitored as gradually reduced and eventually disappeared F_340/380_ fluorescence after the fluorescence reached maximal level (see representative dead cells indicated by arrows) (ns, p>0.05; ***, p < 0.001; ANOVA, Bonferroni’s test; mean ± SEM). (see also Figure S3 for nM2KO versus WT results). (Q-R), Effects of *Trpm2* deletion on mitochondrial function of cortical neurons during OGD evaluated by dequenching of R123 fluorescence. (Q), Representative images of R123-labelled mitochondria before and 30 min after OGD in cultured WT and gM2KO cortical neurons. Control group (no OGD treatment) was used to show the rapid photo bleaching of R123. (R), Average changes of R123 fluorescence 30 min after OGD. WT neurons (n=74 for OGD, n=39 for control) and gM2KO neurons (n=123) were from 4 dishes of cultured neurons isolated from 3 mice (***, p < 0.001; ANOVA, Bonferroni’s test; mean ± SEM). (S-T), Enhanced TRPM2 function by OGD. TRPM2 currents elicited by a ramp protocol ranging from −100 to +100 mV in cultured cortical neurons from WT mice using pipette solutions containing 100 nM Ca^2+^ and 10 μM ADPR (S). Averaged current amplitude (measured at +100 mV) was increased by 1.4-fold after OGD (T) (*, p < 0.05, unpaired *t*-test, mean ± SEM, n=7). (U-V), Up-regulation of TRPM2 by MCAO. TRPM2 expression level analyzed by western blotting (U) was increased by 7.6-fold (V) in the infarcted (right) hemisphere of brains from MCAO WT mice in comparison with sham-operated WT mice 24 hrs after surgery. Protein levels were normalized by β-Tubulin (***, p < 0.001, unpaired *t*-test, mean ± SEM, n=12/group). (W-Z), Deletion of TRPM2 abolishes the increase of surface expression of NMDARs induced by MCAO. (W), Representative western blotting of the surface expression of GluN1a, GluN2a, and GluN2b in cell membrane protein extractions of 3 brains from WT and gM2KO mice subjected to either sham surgery or MCAO. (X-Z), changes of surface expression levels of GluN1a, GluN2a, and GluN2b in WT and gM2KO mice subjected to either sham surgery or MCAO. The entire hemisphere at the operation side (right hemisphere) was harvested 24 hrs after surgery for protein extraction. Protein levels were normalized to the membrane protein loading control pan-cadherin for quantification (ns, no statistical significance, *, p < 0.05, **, p < 0.01, ***, p < 0.001; ANOVA, Bonferroni’s test; mean ± SEM, n=12).

### TRPM2 enhances surface expression of NMDARs and exacerbates excitotoxicity

Various mechanisms are involved in neuronal death, among which Ca^2+^ overload is a major factor. Using cultured cortical neurons, we applied oxygen-glucose deprivation (OGD) conditions to the neurons to mimic *in vivo* ischemic injury, and analyzed changes of intracellular Ca^2+^ and cell death as previously reported (Weilinger et al., 2016). OGD induced a persistent rise of intracellular Ca^2+^ (Figure 1M-O) until the lysis of neurons as reflected by a complete loss of Fura-2 fluorescence. The lysed neurons were counted as dead neurons, as neuronal lysis is a hallmark feature of necrosis (Weilinger et al., 2016). Throughout the 90 mins of OGD perfusion, a noticeable number of neurons (6.8%) in the WT group died after 30 mins of OGD, and neuron death increased to 35.2% and 58.5% at 60 and 90 mins, respectively (Figure 1M-P). In contrast, there was only 1.9%, 8.7%, and 16.3% dead neurons at 30, 60, and 90 mins in the TRPM2-KO group, respectively (Figure 1P), indicating that TRPM2 deletion protects neurons against OGD-induced neuronal death. Similar results were also observed in neurons isolated from neuron-specific *Trpm2* deletion Cre^+^ mice in comparison with Cre^-^ control littermates (Figure S3). Consistent with the higher percentage of neuronal death induced by OGD, the increase in intracellular Ca^2+^ was also remarkably higher in WT than in TRPM2-KO neurons (Figure 1M-O). It was previously shown that during *in vitro* ischemia of cultured cortical neurons, 80% of Ca^2+^ entry is mediated by NMDARs (Goldberg and Choi, 1993; Lipton, 1999). The drastic reduction of Ca^2+^ entry by *Trpm2* deletion (Figure 1M-P) suggests that TRPM2 might have affected NMDAR functions during OGD exposure.

Mitochondrial dysfunction is another hallmark of excitotoxicity and is an early event leading to neuronal death (Keelan et al., 1999; Schinder et al., 1996; Vergun et al., 1999; White and Reynolds, 1996). Dysfunction of mitochondria is characterized by depolarized mitochondria membrane potential caused by opening of the mitochondrial permeability transition pore (mPTP) (Lemasters et al., 2009), which can be monitored by rhodamine 123 (Rh123) fluorescence dequenching assay (Nguyen et al., 1997). OGD-induced mitochondrial depolarization is indicated by increased Rh123 fluorescence in WT neurons, whereas TRPM2 deletion largely prevented mitochondrial depolarization (Figure 1Q-R). Since activation of NMDARs is known to cause mitochondrial depolarization (Abramov and Duchen, 2008; Qiu et al., 2013; Yan et al., 2020), the fact that TRPM2 deletion largely eliminated mitochondrial depolarization during OGD provides another line of evidence suggesting that TRPM2 might have influenced NMDAR function.

Since TRPM2 is sensitive to oxidative stress stimuli, we investigated how TRPM2 is regulated by ischemic stroke *in vivo* and *in vitro*. We used sub-optimal Ca^2+^ and ADPR concentrations (Du et al., 2009a) in the pipette solution for TRPM2 recording in neurons and exposed neurons to OGD. During OGD stimulation, TRPM2 current amplitude was significantly increased in neurons from WT mice (Figure 1S-T), indicating an enhanced channel activity, which might also happen during *in vivo* ischemic stroke. Moreover, TRPM2 expression level in the WT MCAO brains was 7.6-fold higher than in the WT sham control brains (Figure 1U-V).

As both OGD-induced neural death (Figure 1M-P) and mitochondrial dysfunction (Figure 1Q-R) results suggest that TRPM2 might influence NMDAR functions, we evaluated whether TRPM2 influences NMDAR functions in ischemic stroke *in vivo*. Using plasma membrane protein extracts from brains of TRPM2-KO (gM2KO) and WT littermate mice subjected to MCAO or sham procedure, we discovered that the surface expression levels of NMDARs, including GluN1, GluN2a, and GluN2b, were much higher in the WT MCAO mice in comparison with WT sham control mice. Remarkably, the increase in the surface expression level of NMDARs induced by MCAO mice was almost totally abolished by TRPM2 deletion (Figure 1W-Z). This novel finding prompted us to propose that, as an oxidative stress sensor, TRPM2 influences NMDAR surface expression and function during ischemic stroke, thereby exacerbating NMDAR’s excitotoxicity.

### TRPM2 interacts with NMDARs

To understand how TRPM2 may influence NMDAR surface expression in MCAO brains, we tested whether TRPM2 interacts with NMDARs. We first used HEK-293 cells heterologously expressing TRPM2 and the NMDAR subunits GluN1, GluN2a, and GluN2b (GluN1/GluN2a/GluN2b), and performed co-immunoprecipitation (co-IP) experiments. We found that TRPM2 can be pulled down by antibodies specifically against GluN1, GluN2a, and GluN2b, indicating that TRPM2 interacts with the NMDAR protein complex (Figure 2A). In reciprocal co-IP experiments, when TRPM2 was immunoprecipitated by anti-TRPM2, GluN1, GluN2a, and GluN2b were detected in the precipitated complex by western blot (Figure 2B), further indicating that TRPM2 interacts with NMDARs. Since the subunits GluN1, GluN2a, and GluN2b form a heteromeric channel complex, antibodies against any of these subunits may pull down the entire heterotetrameric complex. To determine which subunits interact with TRPM2, we transfected TRPM2 individually with GluN1, GluN2a, or GluN2b. As shown in Figure 2C, TRPM2 interacted with both GluN2a and GluN2b but not GluN1 when they were separately transfected with TRPM2, indicating that in the GluN1/GluN2a/GluN2b complex, GluN2a and GluN2b subunits interact with TRPM2. We then sought to determine whether endogenous TRPM2 interacts with the NMDAR complex. We used brain tissues from WT MCAO mice since TRPM2 is highly up-regulated by MCAO (Figure 1U-V). As shown in Figure 2D-E, GluN1, GluN2a, and GluN2b were able to pull down TRPM2, and reciprocally, anti-TRPM2 was able to pull down GluN1, GluN2a, and GluN2b in the WT MCAO brain lysates but not in the TRPM2-KO MCAO brain lysates. The interaction between TRPM2 and NMDARs in both exogenous expression systems and brains prompted us to determine the functional significance of the interaction.

**Figure 2.**
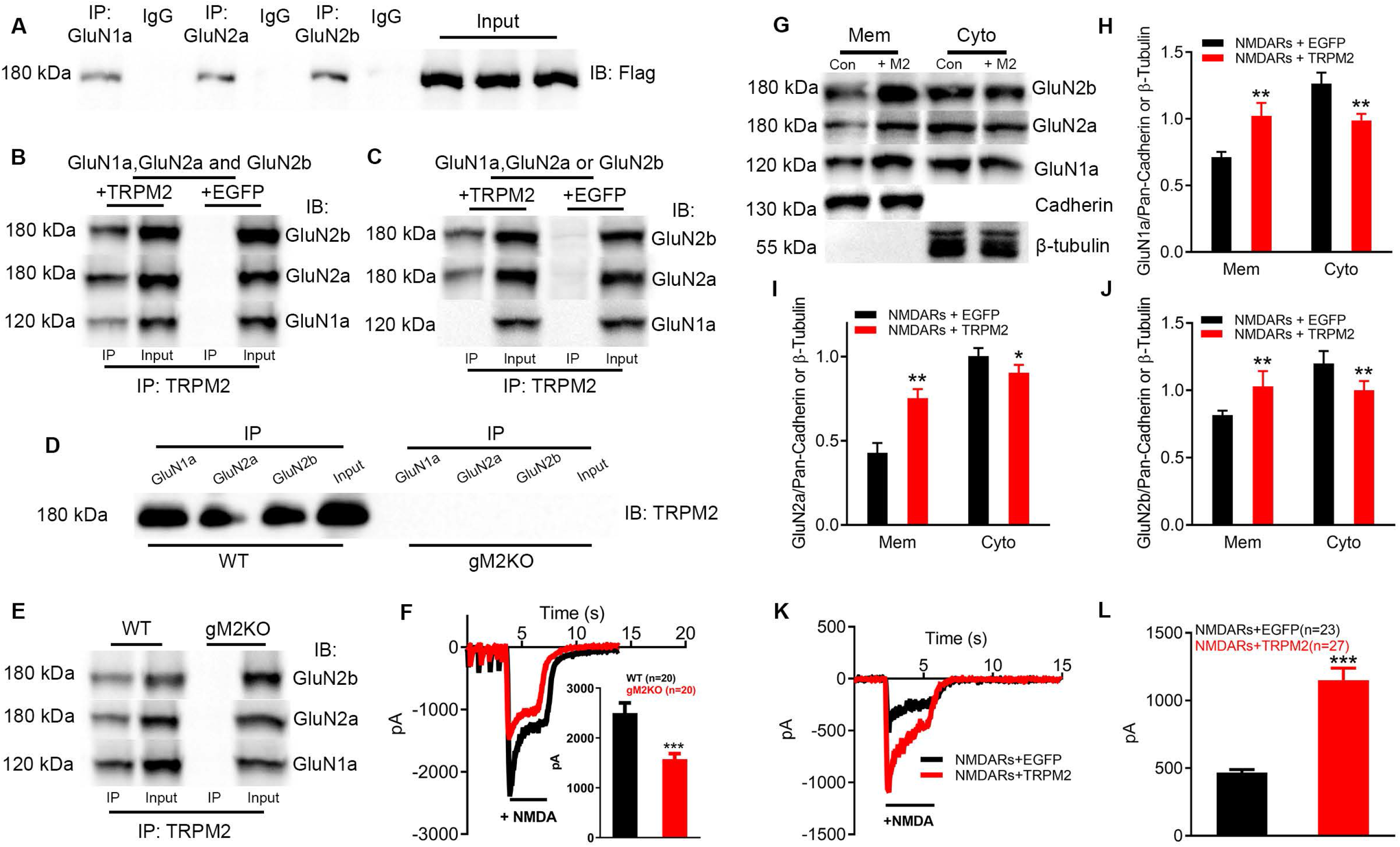
TRPM2 physically and functionally interacts with NMDARs. (A-B), Co-immunoprecipitation (Co-IP) of NMDARs and TRPM2 expressed in HEK-293T cells. NMDAR subunits, GluN1a, GluN2A, and GluN2B were transfected with Flag-tagged TRPM2 (Flag-TRPM2) or EGFP empty vector plasmids. (A), TRPM2 was immunoprecipitated (IP’d) using anti-GluN1a, anti-GluN2a and anti-GluN2b agarose, and detected using western blotting (WB) with anti-Flag. (B), Cell lysates were IP’d using anti-Flag agarose and were probed using WBs with anti-GluN1a, anti-GluN2a, and anti-GluN2b. All the transfections, IP, and WB were replicated 3 times. (C), Co-IP of each subunit of NMDARs and TRPM2 in HEK-293T cells in which Flag-TRPM2 was co-transfected with GluN1a (TRPM2/GluN1a), GluN2a (TRPM2/GluN2a), or GluN2b (TRPM2/GluN2b). EGFP plasmid was used as a control for transfection with each subunit of NMDARs. Cell lysates were IP’d using anti-TRPM2 agarose and were probed using WBs with anti-GluN1a, anti-GluN2a, or anti-GluN2b. TRPM2 interacted with GluN2a and GluN2b, but not GluN1a. All the transfections, Co-IP, and WB were replicated 3 times. (D-E), Endogenous TRPM2 and NMDARs interaction. Co-IP of NMDARs and TRPM2 using protein extractions from brains of WT mice after MCAO. Brain extracts of gM2KO mice subjected to MCAO were used as control. (D) Brain lysates were IP’d using anti-GluN1a, anti-GluN2a, and anti-GluN2b agarose and were probed using WB with anti-TRPM2. (E) Brain lysates were Co-IP’d using anti-TRPM2 antibody and were probed using WBs with GluN1a, GluN2a, and GluN2b. All Co-IP and WB were replicated using at least 3 mouse brains. (F), Representative NMDAR currents recorded by holding at −80 mV in cortical neurons cultured for 14 days from WT and gM2KO mice. NMDA at 10 μM was applied for ∼5 to 10 s. Average peak current amplitude of NMDARs from WT and gM2KO neurons is shown in inset (***, p < 0.001, unpaired *t*-test, mean ± SEM, n=20/group). (G-J), Surface expression changes of NMDARs in HEK-293T cells co-transfected with TRPM2 (NMDARs/TRPM2, or “+M2”), or EGFP plasmid as control (NMDARs/EGFP, or “Con”). (G), Membrane (Mem) and cytosol (Cyto) protein levels assessed with WBs. Pan-cadherin and β-tubulin were used as loading control for membrane and cytosol proteins extracts respectively. (H-J), Quantification of the expression of GluN1a, GluN2a, and GluN2b in cell membrane and cytosol (*, p < 0.05, **, p < 0.01, unpaired *t*-test, mean ± SEM). All transfections, extraction of membrane/cytosolic proteins and immunoblotting were repeated at least 3 times. (K-L), Functional changes of NMDARs when co-expressed with TRPM2 in HEK293T cells. (K), Representative NMDAR currents recorded by holding at −80 mV in HEK293T cells transfected with NMDARs/TRPM2, or NMDARs/EGFP. NMDA at 10 μM was applied for ∼5 to 10 s to activate NMDARs. (L), Average peak current amplitude from NMDARs/EGFP group (n=23) and NMDARs/TRPM2 group (n=27) (***, p < 0.001, unpaired *t*-test, mean ± SEM).

### Functional coupling between NMDARs and TRPM2

We tested NMDAR currents in cultured neurons from WT and TRPM2-KO mice. NMDAR currents elicited by 10 μM NMDA at holding potential of −80 mV were much bigger in WT neurons than that in the TRPM2-KO neurons (Figure 2F). We then tested functional interaction of NMDARs and TRPM2 in the overexpression system. In HEK293T cells overexpressing NMDARs with TRPM2, surface expression levels of GluN1, GluN2a, and GluN2b were higher than that in HEK293T cells overexpressing NMDARs with control EGFP vector plasmids (Figure 2G-J). Consistent with the enhanced surface expression levels, NMDAR currents elicited by NMDA in NMDARs/TRPM2-expressing cells were significantly larger than that in NMDARs/EGFP-expressing cells (Figure 2K-L). The increased surface expression levels and enhanced NMDAR channel functions were also observed in separate transfections when GluN1/GluN2a or GluN1/GluN2b were co-expressed with TRPM2 (Figure S4). These results suggest that TRPM2 and NMDARs physical interactions result in functional coupling reflected by the enhanced functional currents of NMDARs.

### Identification of the NMDAR-interacting domain EE_3_ at the N-terminus of TRPM2

To understand how TRPM2 interacts with NMDARs, we generated N- and C-terminal fragments of TRPM2 tagged with Flag and GFP, respectively (Flag-TRPM2-NT, GFP-TRPM2-CT). When co-expressed with NMDARs, the TRPM2-N but not the TRPM2-C fragment was detected in the precipitate pulled down by GluN1, GluN2a, and GluN2b antibodies (Figure 3A-B). For NMDARs, the C-termini of GluN2a and GluN2b were pulled down by TRPM2, whereas the C-terminal domain-deleted GluN2a (GluN2a-ΔCT) and GluN2b (GluN2b-ΔCT) were absent in the precipitates pulled down by anti-TRPM2 (Figure 3C). These findings suggest that the C-terminal tail of NMARs interact with TRPM2’s N-terminal domain.

**Figure 3.**
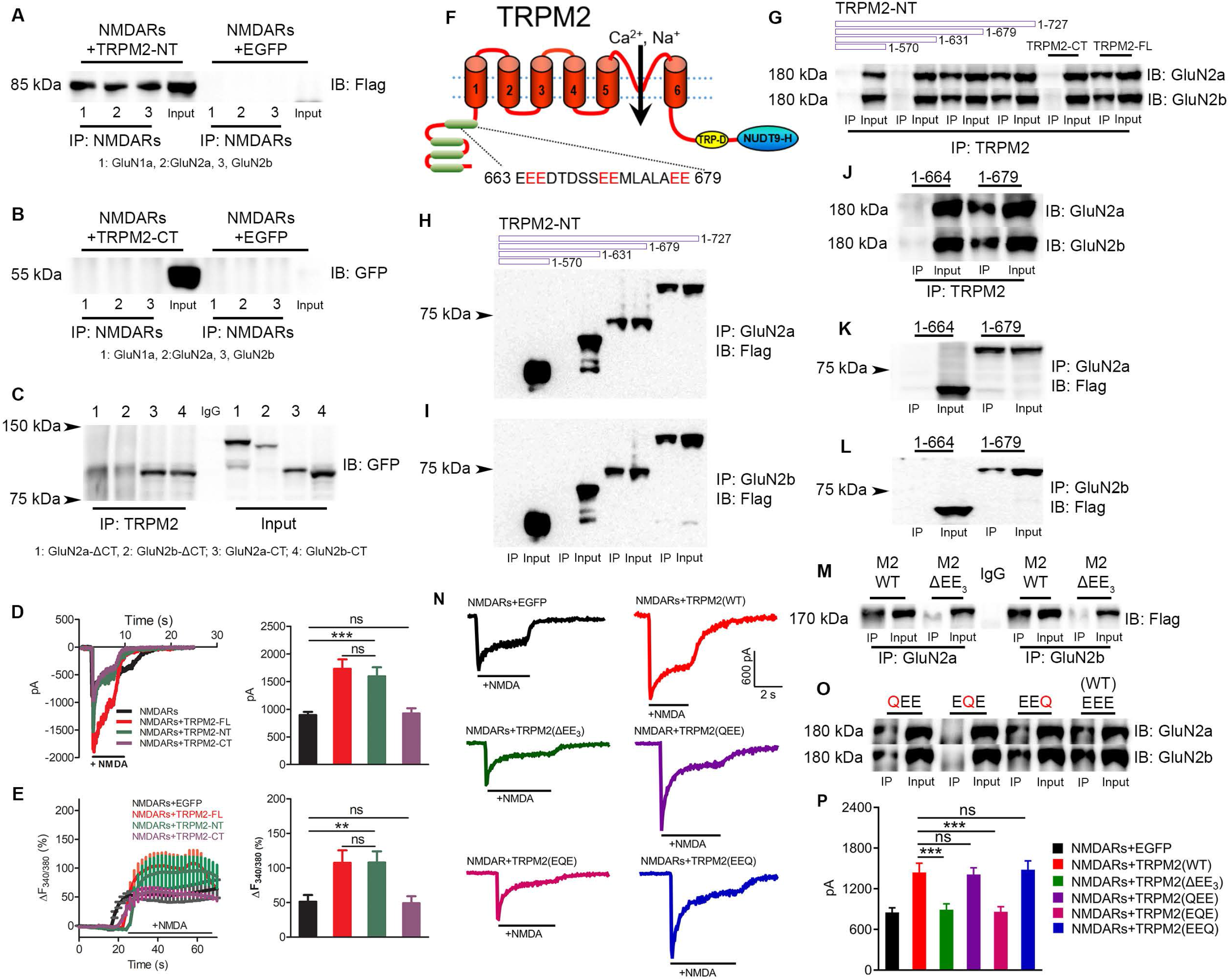
Identification of the interacting domain EE_3_ in the N-terminus of TRPM2 that mediates the physical and functional coupling between TRPM2 and NMDARs. (A-B), Co-IP of N-terminal (1-727) and C-terminal (1060-1503) fragments of TRPM2 (TRPM2-NT, TRPM2-CT) with NMDARs in the HEK293T cells co-transfected with NMDARs or EGFP plasmids. TRPM2-NT (A) and TRPM2-CT (B) were IP’d using anti-GluN1a, anti-GluN2a, or anti-GluN2b antibodies and were probed by WBs using anti-flag (A) or anti-GFP (B). TRPM2-NT was flag-tagged and TRPM2-CT was GFP-tagged. (A), TRPM2-N interacted with NMDARs, as an estimated 85 kDa fragment was detected by anti-flag in NMDARs/TRPM2-N co-transfected cells but not in the NMDARs/EGFP-transfected cells. (B), TRPM2-CT did not interact with NMDARs as the estimated 60 kDa fragment of TRPM2-CT was only detected by anti-GFP in the input of NMDARs/TRPM2-CT co-transfected cell. These results were replicated in three independent experiments. (C), Co-IP of TRPM2 with the C-terminus of GluN2a (GluN2a-CT, 1054-1068), C-terminus deleted GluN2a (GluN2a-ΔC, 1-1053), C-terminus of GluN2b (GluN2b-CT, 1041-1691), and C-terminus deleted GluN2b (GluN2b-ΔC, 1-1047) over-expressed in the HEK293T cells. All the constructs were GFP-tagged. GluN2a-CT, GluN2b-CT, GluN2a-ΔCT, and GluN2b-ΔCT were IP’d using anti-TRPM2 and were probed using WB by anti-GFP. Note that only the GluN2a-CT and GluN2b-CT interacted with TRPM2. These results were replicated in three independent experiments. (D), Representative NMDAR currents recorded in the HEK-293T cells transfected with NMDARs/EGFP, NMDARs/TRPM2-full length (FL), NMDARs/TRPM2-NT, and NMDARs/TRPM2-CT (Left). NMDAR currents were elicited by 10 μM NMDA in tyrode solution perfused for about 7 to 10 s at −80 mV. Average current amplitudes of NMDARs/EGFP group (n=9), NMDARs/TRPM2-FL group (n=8), NMDARs/TRPM2-NT group (n=8), and NMDARs/TRPM2-CT group (n=8) are shown in the bar graph at the right side (ns, p>0.05; ***, p < 0.001; ANOVA, Bonferroni’s test; mean ± SEM). (E), NMDARs-mediated Ca^2+^ influx in HEK-293T cells transfected with NMDARs/EGFP, NMDARs/TRPM2-FL, NMDARs/TRPM2-NT, and NMDARs/TRPM2-CT. Averaged changes of F_340/380_ upon NMDA perfusion at around 15 to 25 s (left), and the mean ΔF_340/380_ measured at ∼40 s is shown in the bar graph (right). (ns, p>0.05; **, p< 0.01; ANOVA, Bonferroni’s test; mean ± SEM; n=17, 14, 13, and 16, respectively, from 3 testing dishes/group). (F), Schematic diagram of the membrane topology of TRPM2. The EE_3_ domain is localized within the MHR4. (G-L), Identification of NMDAR-binding domain at the TRPM2 N-terminus. Co-IP of Flag-TRPM2 N-terminal segments with different lengths (1-570, 1-631, 1-678 and 1-727) with NMDARs co-transfected in HEK-293T cells. (G), lysates were IP’d using anti-TRPM2-N (against TRPM2 N-terminus) and were probed by WBs with anti-GluN2a or anti-GluN2b. TRPM2-CT and TRPM2-FL were included as negative and positive control, respectively. (H, I) Different fragments of Flag-TRPM2 were IP’d with anti-GluN2A (H) or GluN2b (I) and were probed by WBs with anti-Flag. Fragments shorter than 631 aa failed to interact with NMDARs. The results were replicated at least three times. (J-L), Identification of the EE_3_ domain as the binding site for TRPM2 and NMDARs. Co-IP of the TRPM2 N-terminal fragments (1-664 and 1-679) with NMDARs co-expressed in the HEK293T cells. (J), GluN2a and Glu2N2b were IP’d with N-terminal anti-TRPM2 and probed using WBs with anti-GluN2a and GluN2b. (K, L), TRPM2 N terminal segments (1-664 and 1-679) were IP’d with anti-GluN2a (K) and anti-GluN2b, and were probed using WBs with anti-Flag. Fragment 1-664 failed to interact with NMDARs. The critical residues 665-679 for physical interaction together with additional “EE” at the position 680-681 was defined as the EE_3_ binding domain. The results were replicated in three independent experiments. (M-P), Physical and functional coupling of TRPM2 and NMDARs through EE_3_ domain. (M, O), EE_3_ domain deletion mutant of TRPM2 (TRPM2-ΔEE_3_), and EE_3_ mutations of TRPM2, TRPM2-QEE (E666Q and E667Q), TRPM2-EQE (E673Q and E674Q), and TRPM2-EEQ (E680Q and E681Q) were co-expressed with NMDARs in HEK293T cells for co-IP. (M) TRPM2-ΔEE_3_ (M2-ΔEE_3_) was IP’d with GluN2a or GluN2b and probed with anti-Flag. TRPM2-WT (M2-WT) was used as control. (O), TRPM2 mutants QEE, EQE, and EEQ were IP’d with anti-TRPM2 and probed with GluN2a or GluN2b. WT-TRPM2 (EEE) was used as a control. (N), NMDAR currents elicited by NMDA in HEK293T cells co-expressed with empty EGFP vector, WT-TRPM2, TRPM2-ΔEE_3_, TRPM2-QEE, TRPM2-EQE, and TRPM2-EEQ. (P), Mean current amplitude (ns, p<0.05; ***, p < 0.001; ANOVA, Bonferroni’s test; mean ± SEM, n=10/group).

To determine whether the TRPM2-NT fragment is sufficient to cause functional coupling with NMDARs, we co-transfected NMDARs with full-length TRPM2 (TRPM2-FL), TRPM2-NT, or TRPM2-CT plasmids. NMDAR currents recorded in NMDARs/TRPM2-NT expressing cells were similar to those in NMDARs/TRPM2-FL-expressing cells, whereas the currents in NMDARs/TRPM2-CT group were not different from those recorded in cells expressing NMDARs alone (Figure 3D-E), indicating that TRPM2-NT couples with NMDARs.

To further narrow down the NMDAR-interacting domain at the N-terminus of TRPM2 (Figure 3F), we generated a series of N-terminal truncation constructs by incrementally deleting about 50 residues, and tested which fragments interact with NMDARs (Figure 3G-I). We found that the N-terminal amino acid residues from 631 to 679 are critical for the TRPM2 interaction with NMDARs (Figure 3G-I), as both forward co-IP and reverse co-IP confirmed the interaction for the fragments of 1-727 and 1-679, whereas the fragments shorter than 631 residues failed to interact with NMDARs. We further deleted the fragment 1-679 and found that the amino acid resides between 665 and 679 are essential for the interaction of TRPM2 and NMDARs (Figure 3J-L). Interestingly, the 15 residues between 665 and 679 contain two “glutamate-glutamate” (EE) repeats separated by five residues and followed by another EE repeat (Figure 3F), so we named the residues between 665 to 681 the “EE_3_” domain for simplicity. This EE_3_ domain is present in TRPM2 from different species, but not present in other TRPM channels (Figure S5). When the EE_3_ domain was deleted from the full-length TRPM2 (TRPM2-ΔEE3), interaction between TRPM2 and NMDARs was completely disrupted (Figure 3M). Intriguingly, when the middle “EE” of the EE_3_ domain was replaced by “QQ” (glutamine), the TRPM2-EQE mutant failed to interact with NMDARs, whereas mutations of the first and third EE repeats (TRPM2-QEE, TRPM2-EEQ) did not influence the TRPM2-NMDARs interaction (Figure 3O). These results indicate that the EE residues in the middle of the EE3 domain are critical for the TRPM2-NMDAR interaction.

To investigate the functional consequence of disrupting the physical interaction between TRPM2 and NMDARs, wild-type TRPM2 or TRPM2 EE_3_ mutants were co-expressed with NMDARs in HEK293 cells for current recording. Consistent with the disrupted interaction, TRPM2-ΔEE3 and TRPM2-EQE mutants failed to enhance NMDAR currents, whereas TRPM2-QEE and TRPM2-EEQ mutants increased NMDAR currents, similar to the potentiation of NMDAR currents induced by WT-TRPM2 when co-expressed with NMDARs in HEK-293T cells (Figure 3N & 3P). It is noteworthy that the TRPM2 mutations did not affect TRPM2 channel function (Figure S5). These results indicate that the EE_3_ domain is essential for physical and functional coupling between TRPM2 and NMDARs.

### Mechanisms of TRPM2 and NMDARs functional coupling

The above results indicate that TRPM2 interacts with NMDARs through the TRPM2-EE_3_ domain thereby enhancing NMDAR function by increasing its surface localization (Figure 2 & 3). Next, we asked how the interaction of TRPM2 with NMDARs enhances surface expression of NMDARs. Since it has been previously shown that PKC regulates NMDAR trafficking to the cell surface (Lan et al., 2001; Zheng et al., 1999), we reasoned that PKC could be part of the NMDAR-TRPM2 complex. Thus, we investigated whether PKC interacts with TRPM2. Using WT and global TRPM2-KO (gM2KO) mouse brains after MCAO, we found that neuron-specific PKCγ can be readily pulled down by anti-TRPM2 in WT but not in TRPM2-KO (gM2KO) brains (Figure 4A). In HEK293 cells over-expressing PKCγ with full-length TRPM2 (TRPM2-FL), TRPM2-CT, or TRPM2-NT, we found that TRPM2-FL and TRPM2-NT, but not TRPM2-CT, pulled down PKCγ, indicating that PKCγ interacts with TRPM2-NT (Figure 4B). Moreover, when we used brain tissues from both sham and MCAO mice, the amount of PKCγ pulled down by anti-TRPM2 antibody in MCAO mice was about 1.5-fold of that from sham mice (Figure 4C-D), suggesting that oxidative stress during MCAO increased TRPM2 and PKCγ interaction. Indeed, cultured neurons treated with H_2_O_2_ exhibited significantly increased TRPM2 and PKCγ interaction (Figure 4E-F).

**Figure 4.**
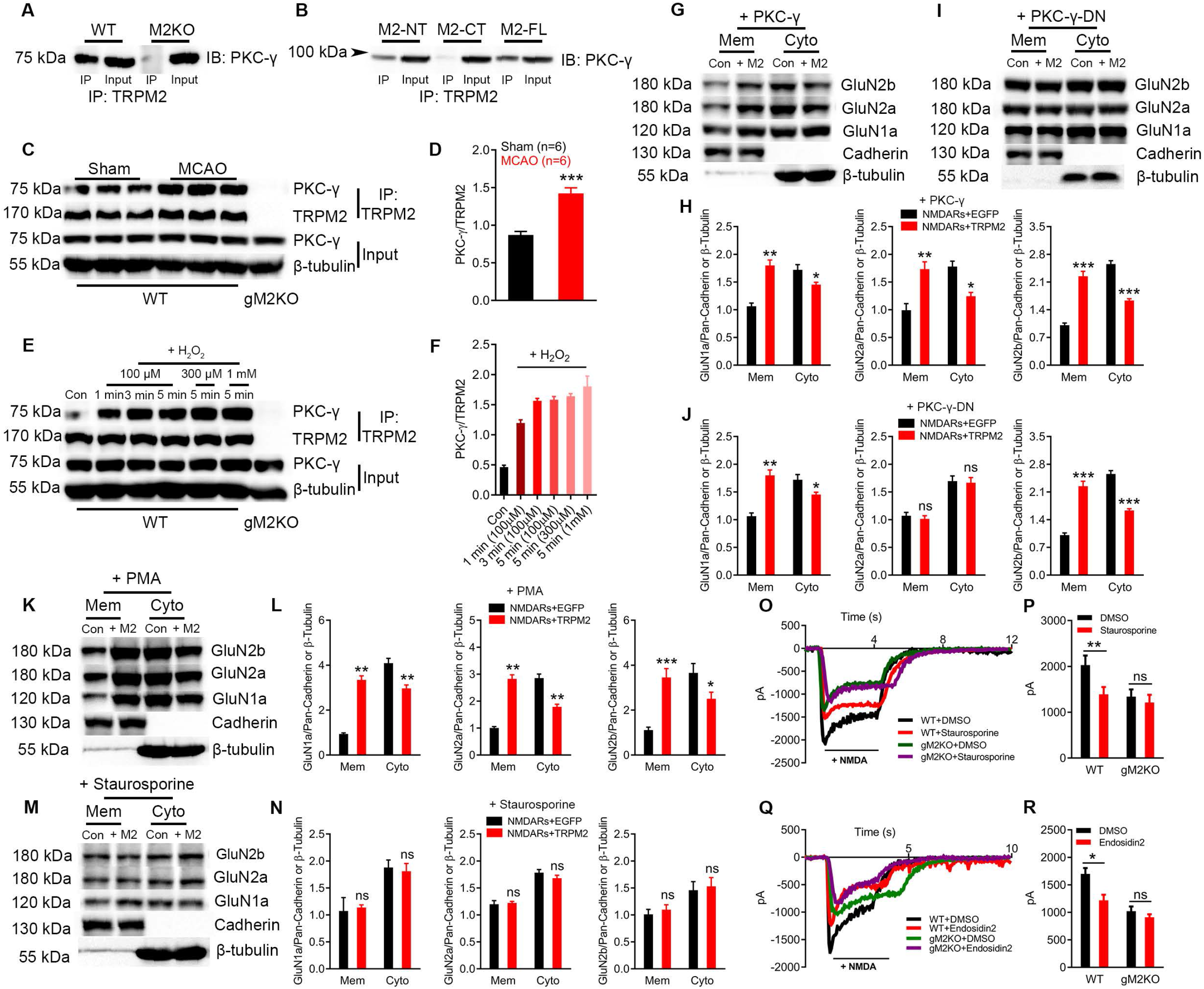
N-terminal domain of TRPM2 Interacts with PKC-γ. (A-B), Co-IP of PKCγ and TRPM2 using proteins extracted from mouse brains after MCAO (A), and in HEK293T cells expressing PKCγ with TRPM2-FL, TRPM2-NT, or TRPM2-CT (B). PKC-γ was IP’d with anti-TRPM2 and was probed by WBs using anti-PKCγ. The results were replicated in 3 independent experiments. (C,D), Oxidative stress promotes TRPM2 and PKCγ interaction in MCAO brains. (C), Co-IP of TRPM2 and PKCγ using protein extracts from WT mice brains subjected to MCAO or sham operation. PKC-γ was IP’d using anti-TRPM2 and was probed with anti-PKCγ by WB. TRPM2 was used as control for IP protein, and β-tubulin was used as loading control. (D) Mean PKCγ/TRPM2 ratios in sham and MCAO mice. (***, p < 0.001, unpaired *t*-test, mean ± SEM; n=6). (E,F), Effects of H_2_O_2_ on TRPM2 and PKCγ interaction in cultured neurons. (E), PKCγ was IP’d with anti-TRPM2 in protein extracts from cultured WT cortical neurons exposed to H_2_O_2_ (at 100 µM for 1 min, 3 min, and 5 min, at 300 µM for 5 min and at 1 mM for 5 min), and probed with anti-PKC-γ. For IP proteins, TRPM2 was used as loading control. For lysates, β-tubulin was used as loading control. (F), Average PKCγ normalized to TRPM2 from 3 independent experiments. (***, p < 0.001; ANOVA, Bonferroni’s test; mean ± SEM, n=3/group). (G-J), Surface expression of NMDARs in HEK-293T cells co-transfected with PKC-γ/EGFP (Con) and PKC-γ/TRPM2 (+M2) (G) or PKC-γ-DN/EGFP (Con) and PKC-γ-DN/TRPM2 (+M2) (I). Membrane and cytosol protein levels of NMDARs were quantified from 3 independent experiments (H, J). (*, p < 0.05, **, p < 0.01, ***, p < 0.001, unpaired *t*-test, mean ± SEM). (K-N), Surface expression of NMDARs in HEK-293T cells co-transfected with TRPM2 or EGFP with the treatment of PKC activator PMA (K) or inhibitor Staurosporine (M). Membrane (Mem) and cytosol (Cyto) NMDARs were quantified from 3 independent experiments (L, N). (*, p < 0.05, **, p < 0.01, ***, p < 0.001, unpaired *t*-test, mean ± SEM, n=3/group). (O,P), Effects of staurosporine on NMDAR currents recorded from cortical neurons cultured for 14 days. (O), Representative NMDAR currents elicited at −80 mV by 10 μM NMDA in WT and TRPM2-KO (gM2KO) neurons treated with or without Staurosporine at 1 µM for overnight. (P), Mean current amplitude (ns, p>0.05; **, p < 0.01; ANOVA, Bonferroni’s test; mean ± SEM, n=11, 10, 10, and 12 neurons from 2 mice, respectively). (Q-T), Effects of Endosidin2 on NMDA currents recorded from cortical neurons cultured for 14 days. (S) Representative NMDAR currents elicited by NMDA in WT and TRPM2-KO neurons treated with or without endosidin2 at 1 μM overnight. (T) Average current amplitude (ns, p>0.05; *, p < 0.05; ANOVA, Bonferroni’s test; mean ± SEM, n=10 neurons/group from 2 mice respectively).

In order to investigate whether PKCγ influences surface expression levels of NMDARs, we co-expressed NMDARs and TRPM2 with wild-type PKCγ or dominant-negative PKCγ (PKCγ-DN). As shown in Figure 4G-J, over-expression of PKCγ further increased surface expression of NMDARs, whereas dominate-negative PKCγ abolished the increase of surface NMDAs by TRPM2, suggesting that a functional PKCγ is required for increased NMDARs trafficking induced by TRPM2.

We also used pharmacological tools to probe the function of PKC in the TRPM2-mediated enhancement of surface NMDAR levels. HEK293T cells transfected with NMDARs and TRPM2 exhibited much higher level of surface NMDARs after treatment with PKC activator PMA (Figure 4K-L), whereas PKC inhibitor staurosporine abolished the enhanced surface expression level of NMDARs (Figure 4M-N). The effects of PKC on NMDARs were further confirmed by the effects of PKC inhibitor staurosporine on NMDAR currents recorded in the neurons from WT mice. Staurosporine normalized NMDAR current amplitude in the WT neuron to the similar amplitude level of NMDAR current in TRPM2-KO neurons (Figure 4O-P).

As trafficking of NMDARs to plasma membrane involves exocysts (Sans et al., 2003), we tested the effects of blocking exocysts on NMDARs surface expression. Endosidin 2, an exocyst inhibitor, prevented the increased surface expression level of NMDARs (Figure S6) induced by co-expression with TRPM2, and abolished the increased current amplitude of NMDARs by TRPM2 in WT neurons (Figure 4Q-R).

Taken together, the above results indicate that PKCγ interacts with TRPM2, which can be promoted by an oxidative stress condition and by MCAO *in vivo*, and that interaction of PKCγ with TRPM2 enhances NMDARs surface trafficking likely via exocysts.

### Disrupting peptide eliminates physical interaction and functional coupling of TRPM2 and NMDARs

As the TRPM2 N-terminal EE_3_ domain is critical for TRPM2 and NMDARs interaction, we designed membrane-permeable peptides, TAT-EE_3_ and scrambled control peptide TAT-SC, to investigate whether disruption of physical interaction eliminates the functional coupling between TRPM2 and NMDARs. To determine whether TAT-EE_3_ is able to disrupt the interaction of TRPM2 and NMDARs, we conducted co-IP experiments using NMDARs co-expressed with WT-TRPM2 and TRPM2 EE_3_ mutants including TRPM2-QEE, TRPM2-EQE, TRPM2-EEQ, and the deletion mutant TRPM2-ΔEE_3_. Similar to the TRPM2-Δ(EE)_3_ and TRPM2-EQE mutants, TAT-EE_3_ treatment completely disrupted the interaction of TRPM2 with GluN2a (Figure 5A) and GluN2b (Figure 5B), whereas TAC-SC treatment and mutants TRPM2-QEE and TRPM2-EEQ did not influence TRPM2 and GluN2a and GluN2b interactions. We further determined the surface expression of NMDARs after TRPM2- and NMDAR-overexpressing cells were treated with TAT-EE_3_ or TAT-SC. TAT-EE_3_ eliminated the enhancement of NMDAR surface expression by TRPM2 (Figure 5D, F), whereas TAT-SC exhibited no influence (Figure 5C, E). Consistent with the disrupted interaction between TRPM2 and NMDARs and the eliminated increase of NMDAR surface expression, the NMDAR current amplitude in NMDARs and TRPM2-overexpressing cells treated with TAT-EE_3_ was significantly smaller than that in cells treated with TAT-SC (Figure 5G-H).

**Figure 5.**
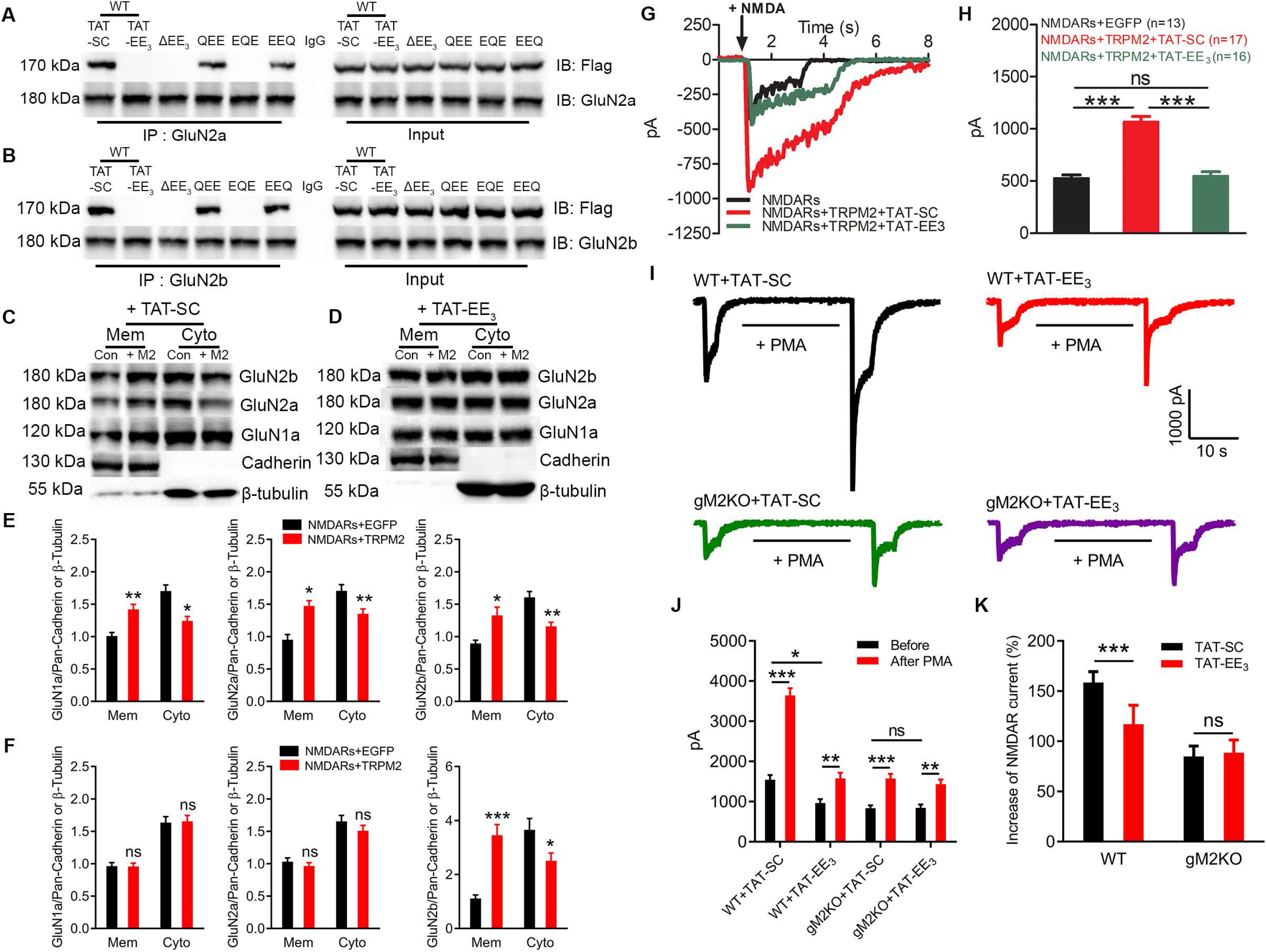
TAT-EE_3_ disrupts the physical and functional interaction between TRPM2 and NMDAR. (A-B), Effects of TAT-EE_3_ on the interaction of TRPM2 and NMDARs. Co-IP tests of NMDARs with WT or EE_3_-domain mutants of TRPM2 (Flag-tagged) in co-transfected HEK293T cells. WT-TRPM2 groups were treated with 10 µM TAT-SC or TAT-EE_3_ overnight. Lysates were IP’d with anti-GluN2a (A) or anti-GluN2b (B), and probed by WBs with anti-Flag. IgG was used as control for IP. Similar to the mutants TRPM2-ΔEE_3_ and TRPM2-EQE, TAT-EE_3_ disrupted TRPM2 interactions with GluN2a and GluN2b. These results were replicated in 3 independent experiments. (C-F), Effects of TAT-EE_3_ on surface expression of NMDARs. HEK293T cells co-transfected with NMDARs and TRPM2 (+M2) or EGFP vector (Con) were treated with 10 µM TAT-SC (C) or TAT-EE_3_ (D) overnight. Membrane (Mem) and cytosol (Cyto) proteins of NMDARs in TAT-SC (E) and TAT-EE_3_ (F) treated groups were assessed by WBs and analyzed in reference to Pan-cadherin or β-tubulin (**, p < 0.01, ***, p < 0.001, unpaired *t*-test, mean ± SEM). All the results were replicated in at least 3 independent experiments. (G-H), Effects of TAT-EE_3_ on NMDAR currents in HEK-293T cells transfected with NMDARs and TRPM2. Transfected cells were treated with 10 μM TAT-EE_3_ or TAT-SC for overnight. (G), Representative NMDAR currents elicited at −80 mV by exposing to 10 μM NMDA for about 2 to 4 s. (H), Mean current amplitude of NMDARs/EGFP (n=13), and NMDARs/TRPM2 treated with TAT-SC (n=17) or TAT-EE3 (n=16) groups (ns, p>0.05; ***, p < 0.001; ANOVA, Bonferroni’s test; mean ± SEM). (I-K), Effects of TAT-EE_3_ on PKC-induced changes of NMDAR currents recorded in cortical neurons of WT and TRPM2-KO. Neurons were incubated with 10 μM TAT-SC or TAT-EE_3_ overnight before current recording. (I) Representative NMDAR currents elicited by 10 μM NMDA at −80 mV before and after PMA (1 μM) perfusion for 20 s. TAT-SC or TAT-EE3 10 µM was included in the pipette solution for current recording. (J), Average NMDAR current amplitude before and after 1 μM PMA from WT and TRPM2-KO neurons treated with TAT-SC or TAT-EE_3_. (K), Average percentage increases of NMDAR currents induced by PMA in WT and TRPM2-KO neurons treated with TAT-SC or TAT-EE_3_ (ns, *p>0.05; ***, p < 0.001; ANOVA, Bonferroni’s test; mean ± SEM, n=19, 14, 14 and 11, respectively).

As PKC activation can both increase NMDAR trafficking to the cell surface and regulate channel activity (Lan et ^a^l., 200^1^), we sought to determine the PMA effects on NMDAR currents in neurons treated with TAT-EE_3_ or TAT-SC. Cultured neurons from WT and TRPM2-KO mice were treated with TAT-EE_3_ or TAT-SC overnight. NMDAR currents were elicited before and after 20 s perfusion with PMA. As shown in Figure 5I (top left), NMDA-induced current was increased from 1542.1±117.1 pA to 3642.1±180.8 pA (Figure 5J) by PMA perfusion for 20 s (Figure 5J-K), about a 1.5-fold increase, in neurons from WT mice. However, in the neurons incubated with TAT-EE_3_, the NMDAR current amplitude was much smaller before and after PMA perfusion (Figure 5I, top right and Figure 5J), and the increase of current amplitude induced by PMA was also significantly smaller than that in neurons incubated with TAT-SC (Figure 5K). The smaller NMDAR currents before PMA and smaller increase of NMDAR currents after PMA in TAT-EE_3_-incubated neurons suggest that disruption of TRPM2 and NMDARs interaction may have affected surface trafficking of NMDAR-induced PKC activation. In the neurons from global TRPM2-KO (gM2KO) mice, the NMDAR current amplitude (Figure 5I, bottom) and PMA-induced increase in NMDAR currents were much smaller than that of WT neurons. Moreover, there was no difference in PMA-induced changes of NMDAR currents between TAT-SC and TAT-EE_3_ incubated neurons from TRPM2-KO mice (Figure 5J-K). These results indicate that similar to TRPM2-KO, disruption of the interaction between TRPM2 and NMDARs in the WT neurons by TAT-EE_3_ reduced NMDAR currents, and attenuated the PMA-induced increase in NMDAR currents. As PMA causes an increase of NMDAR channel activities and surface trafficking, it is conceivable that in TRPM2-KO and TAT-EE_3_-treated neurons, PMA failed to cause surface trafficking of NMDARs. This notion is further supported by the fact that inhibition of exocysts abolished the increase of surface expression of NMDARs co-expressed with TRPM2 in HEK293T cells (Figure S6), and the enhancement of NMDAR currents by TRPM2 in WT neurons (Figure 4Q-R).

### Disruption of TRPM2 and NMDAR interaction protects neurons against ischemic injury *in vitro*

The functional uncoupling of TRPM2 from NMDARs by TAT-EE_3_ prompted us to investigate its significance during ischemic injury. Using cultured neurons, the disrupting peptide TAT-EE_3_ efficiently reduced NMDAR currents in neurons from WT mice but not from TRPM2-KO mice, whereas TAT-SC exhibited no influence on NMDAR currents from WT and TRPM2-KO neurons (Figure 6A-B), indicating the specificity of TAT-EE_3_. To mimic *in vivo* ischemia conditions, we used OGD to treat cultured neurons. When neurons were exposed to OGD, the changes of intracellular Ca^2+^ were significantly reduced in WT but not in TRPM2-KO neurons by pretreatment with TAT-EE_3_ (Figure 6C-D). The OGD-mediated neuronal death was 10.6%, 47.6%, and 70.5% at 30, 60, and 90 min, respectively, in WT neurons, which was drastically reduced to 3.6%, 9.9%, and 45.2% by TAT-EE_3_, similar to the percentage death rate in the global TRPM2-KO (gM2KO) neurons (Figure 6E).

**Figure 6.**
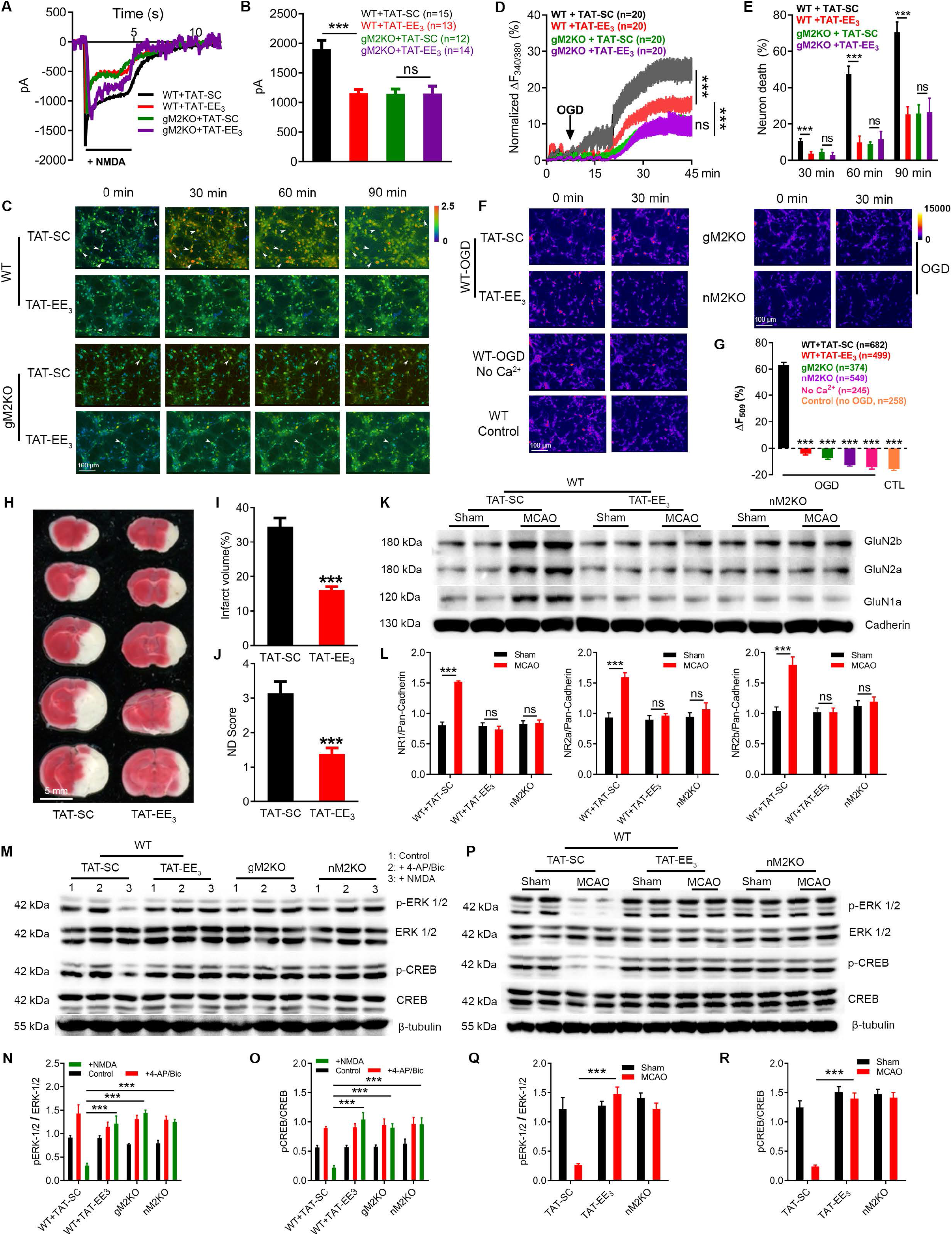
Uncoupling TRPM2 and NMDARs by TAT-EE_3_ protects neurons against ischemic injury *in vitro* and *in vivo* by preserving pro-survival signaling. (A-B), Effects of TAT-EE_3_ on NMDAR currents in WT and gM2KO cortical neurons cultured for 14 days. (A), Representative recording of NMDAR currents in cultured neurons treated with TAT-SC or TAT-EE3 at 10 µM overnight before current recording. TAT-SC or TAT-EE3 10 μM was included in the pipette solution during current recording. (B), Average current amplitude in WT and gM2KO neurons treated with TAT-SC or TAT-EE_3_ (ns, p>0.05; ***, p < 0.001; ANOVA, Bonferroni’s test; mean ± SEM, n=15, 13, 12, and 14, respectively). (C-E), Effects of TAT-EE_3_ on Ca^2+^ overload and neuronal death caused by OGD. Cultured WT and gM2KO cortical neurons were treated with 10 μM TAT-SC and TAT-EE3 overnight before experiments. (C), Ratio Ca^2+^ imaging showing intracellular Ca^2+^ changes induced by OGD in TAT-EE_3_- or TAT-SC-treated WT and gM2KO neurons. Fluorescence ratio F_340/380_ was used to represent intracellular Ca^2+^ changes (see the scale bar at the right shoulder of panel C). Neurons with increasingly elevated Ca^2+^ levels such as the ones indicated with arrows die with time, as reflected by the disappearance of the fluorescence at the next time point. Ionomycin (Iono) was used at the end of the experiments to serve as an internal control for normalizing F_340/380_ (not shown). (D), Ca^2+^ increases represented by normalized F_340/380_ induced by OGD in the first 45 min analyzed from 20 neurons randomly chosen from each group. (E), Average neuronal death rate (see panel (C)) evaluated at 30, 60, and 90 min after OGD. Neuronal death was monitored as F_340/380_ fluorescence gradually reduced and eventually disappeared after the fluorescence reached maximal level (see representative dead cells indicated by arrows in (C)) (ns, p>0.05; ***, p < 0.001; ANOVA, Bonferroni’s test; mean ± SEM). (F-G), Effects of TAT-EE_3_ on mitochondrial function evaluated using R123 de-quenching assay. Cultured cortical neurons from gM2KO mice, nM2KO mice, and WT control littermates were incubated with 10 µM TAT-SC or TAT-EE3 for overnight before OGD exposure. (F), R123 fluorescence changes in neurons of different groups induced by OGD. External perfusion solution aCSF containing no Ca^2+^ (No Ca^2+^) was used as a control. WT neurons without OGD exposure were used as a control (WT control) to illustrate the rapid photo bleaching of Rh123. (G), Mean R123 fluorescence changes induced by 30 min OGD exposure in TAT-SC (n=682), TAT-EE_3_ (n=499), Global M2KO (n=374), neuronal M2KO (549) neurons, or no-OGD Control (=258). The accumulated numbers of neurons in each group for data analysis were from 3∼5 independent experiments using neurons isolated from 3 mice/group (***, p < 0.001; ANOVA, Bonferroni’s test; mean ± SEM). (H-J), TAT-EE_3_ reduces infarct size and improves neurological deficit (ND) score in WT MCAO mice. (H), TTC staining of brain slices 24 hrs after MCAO from WT mice intraperitoneally (ip) administrated with TAT-SC or TAT-EE_3_ at 100 nmol/kg 15 min before MCAO procedure. (i), Mean infarct volume after MCAO. (J), Average neurological deficit score. (***, p < 0.001; ANOVA, Bonferroni’s test; mean ± SEM; n=7 for TAT-SC and n=8 for TAT-EE_3_ groups). (K-L), Effects of TAT-EE_3_ on surface expression of NMDARs in WT and nM2KO mice 24 hrs after MCAO or sham procedure. TAT-SC and TAT-EE_3_ were ip administrated (100 nmol/kg) 15 mins before MCAO or sham procedure. (K) WB of surface expression of NMDARs from two representative samples/group. Pan-cadherin was used as loading control. (I), Quantification of surface expression of NMDARs normalized to Pan-cadherin (ns, p>0.05; ***, p < 0.001; ANOVA, Bonferroni’s test; mean ± SEM; n=4 mice/group). (M-O), Effects of TAT-EE_3_ on ERK1/2 and CREB activities in cultured neurons isolated from gM2KO and nM2KO mice, and WT littermates. WT neurons were treated with TAT-EE_3_ or TAT-SC overnight before the experiments. Neurons were incubated for 1 h before experiments with 1) control (PBS), 2) NMDA (10 μM) to activate synaptic and extrasynaptic NMDARs, and 3) 4-AP 2.5 mM plus bicuculine (Bic) 50 μM (4-AP/Bic) to activate synaptic NMDARs. (M), ERK1/2 and CREB activities were evaluated by detecting phosphorylated ERK1/2 (p^44/42^ERK1/2) and CREB (pSer^133^CREB). (N-O), Quantification of the active CERB and ERK1/2 in each group under three different treatment conditions (***, p < 0.001; ANOVA, Bonferroni’s test; mean ± SEM, n=3). (Q-R), Effects of TAT-EE_3_ and *Trpm2* deletion on ERK and CREB activities in mice subjected to MCAO. (P), ERK and CREB activities were evaluated by assessing the levels of pERK1/2 and pCREB in nM2KO mice and WT control littermates subjected to MCAO or sham surgery. TAT-EE_3_ or TAT-SC was ip administrated to WT mice 15 min before MCAO or sham surgeries. (Q, R), Quantification of pERK1/2 and pCREB expressed in the brains after MCAO or sham procedure (***, p < 0.001; ANOVA, Bonferroni’s test; mean ± SEM, n=4/group).

We next evaluated whether TAT-EE_3_ influences OGD-mediated mitochondrial dysfunction using the Rh123 fluorescence de-quenching assay (Nguyen et al., 1997). As shown in Figure 6F-G, OGD caused mitochondria depolarization as reflected by the increase of Rh123 fluorescence in WT neurons treated with TAT-SC. In contrast, mitochondrial membrane depolarization induced by OGD was markedly inhibited in WT neurons treated with TAT-EE_3_, similar to the inhibited depolarization in neurons from global TRPM2-KO (gM2KO) or neuron-specific TRPM2 deletion (nM2KO) mice (Figure 6F-G). The mitochondria membrane depolarization was mediated by Ca^2+^ entry because when neurons were perfused with Ca^2+^ free OGD, membrane depolarization was inhibited (Figure 6F: micrographs at 2^nd^ row from bottom, and Figure 6G). These results indicate that the TAT-EE_3_ disrupting peptide abolishes the exacerbation of NMDAR excitotoxicity that results from functional coupling of TRPM2 and NMDARs, and protects neurons from ischemic injury *in vitro*.

### Disruption of TRPM2 and NMDAR interaction protects mice against ischemic stroke

To determine if TAT-EE_3_ inhibits ischemic injury *in vivo*, we administrated TAT-EE_3_ or TAT-SC (100 nmol/kg) 15 mins before MCAO or sham surgery as previously reported (Weilinger et al., 2016). Infarct volume and neurological deficit scores were evaluated 24 hrs after MCAO (see description for Figure 1). As shown in Figure 6H-J, TAT-TTE_3_-treated mice exhibited significantly reduced infarct volume and improved neurological behavior scores in comparison with TAT-SC-treated mice. Similar to those in the global (gM2KO) (Figure 1W) and neuronal (nM2KO) TRPM2-KO mice (Figure 6K), the enhanced surface expression of NMDARs was abolished by TAT-EE_3_ (Figure 6K-L) in MCAO mice, indicating that disruption of NMDARs and TRPM2 coupling is an effective approach to protect the brain against ischemia injury.

As synaptic NMDARs promote pro-survival signals whereas extrasynaptic NMDARs promote pro-death signals (Hardingham and Bading, 2010; Hardingham et al., 2002), we sought to determine whether the enhanced NMDAR surface expression by MCAO in WT mice is pro-survival or pro-death. Cultured neurons from global TRPM2-KO, neuronal TRPM2-KO (Cre^+^), or Cre^-^ WT littermates (WT) were treated for 1 h with NMDA to activate both synaptic and extra-synaptic NMDARs, or 4-AP (2.5 mM) plus 50 μM bicuculline (Bic) to activate only synaptic NMDARs (Hardingham et al., 2002; Nicolai et al., 2010). WT neurons were pre-incubated with TAT-SC or TAT-EE_3_ overnight. After induction of NMDAR activation by NMDA or 4-AP/Bic, neurons were harvested for quantitative analysis of CREB and ERK1/2 activation. In neurons pre-incubated with TAT-SC, treatment with 4-AP/Bic significantly increased neuronal survival signal, as indicated by the phosphorylated ERK1/2 (pERK1/2) and phosphorylated CREK (pCREB) in comparison with control group, whereas NMDA treatment drastically reduced the pro-survival signal pERK1/2 and pCREB in WT neurons treated with TAT-SC group (Figure 6M-O), similar to previous findings in neurons without any pre-incubation (Hardingham et al., 2002; Nicolai et al., 2010). In stark contrast, both 4-AP/Bic and NMDA treatments increased pro-survival signal (pERK1/2 and pCREB) levels in WT neurons pre-incubated with TAT-EE_3_, as well as in neurons from mice with global or neuronal-specific TRPM2 deletion, indicating that disruption of the TRPM2-NMDAR interaction by TAT-EE_3_ or by deletion of TRPM2 largely eliminated extrasynaptic activation of NMDARs by NMDA.

To determine whether TRPM2 exacerbates NMDAR extrasynaptic excitotoxicity in WT MCAO mice, we further analyzed CREB and ERK1/2 activities in brain tissues from MCAO mice treated with TAT-SC or TAT-EE3. In TAT-SC-treated mice, pERK1/2 and pCREB levels were significantly lower in MCAO brains than sham control brains, whereas MCAO induced no reduction in pERK1/2 and pCREB levels in TAT-EE_3_-treated brains (Figure 6P-R). Similarly, neuronal TRPM2-KO (Figure 6P-R) and global TRPM2-KO (Figure S7A-C) prevented reduction of pERK1/2 and pCREB levels after MCAO, indicating that *in vivo* disruption of TRPM2-NMDAR interaction, as well as deletion of TRPM2, protects brain against ischemia injury likely by inhibiting the extrasynaptic NMDAR-induced large reduction in pro-survival signals.

## DISCUSSION

In this study, we report a previously unknown mechanism underlying ischemic brain stroke. We discovered that the up-regulation of TRPM2 during ischemic stroke is physically and functionally coupled to NMDARs, which results in enhanced extrasynaptic NMDAR activity and thereby leads to increased excitotoxicity. We revealed that the mechanism by which TRPM2 increases extrasynaptic NMDAR excitotoxicity is through interaction between TRPM2 and PKCγ, which leads to increased NMDAR protein trafficking and surface expression. Moreover, we identified the specific binding domain in the N-terminus of TRPM2 for coupling of TRPM2 with NMDARs, and designed a membrane-permeable disrupting peptide to uncouple TRPM2 from NMDARs. By disrupting the TRPM2-NMDAR interaction, the peptide TAT-EE_3_ not only inhibited OGD-induced excitotoxicity *in vitro*, but also efficiently protected mice against ischemic stroke injury *in vivo*. Our results indicate that by exacerbating the excitotoxicity of NMDARs, TRPM2 converges both excitotoxic and non-excitotoxic Ca^2+^ signaling pathways in mediating neuronal death in ischemic stroke. This new mechanism may serve as a foundation for designing and developing effective strategies for future ischemic stroke therapies.

### TRPM2 in neurons plays a key role in ischemic injury by exacerbating NMDARs excitotoxicity

The Ca^2+^-permeable TRPM2 was discovered as an oxidative stress-activated non-selective cation channel (Hara et al., 2002; Perraud et al., 2001; Sano et al., 2001). Although TRPM2 has been implicated in ischemic stroke (Belrose and Jackson, 2018), the mechanisms by which TRPM2 mediates ischemic injury have been controversial. Some studies suggest that neuronal TRPM2 contributes to ischemic injury (Alim et al., 2013; Jia et al., 2011), whereas others report that TRPM2 expressed in immune cells contributes (Gelderblom et al., 2014). As all the previous studies used global TRPM2-KO (gM2KO), we established neuron-specific TRPM2 knockout (nM2KO) and found that neuronal TRPM2 plays a key role in mediating ischemic brain injury. Our results not only establish that TRPM2 expressed in neurons is critical for mediating ischemic injury, but also reveal a previously unknown mechanism that TRPM2 exacerbates NMDAR excitotoxicity in mediating neuronal death during ischemic injury.

### A specific domain mediates physical and functional coupling of TRPM2 with NMDARs and exacerbation of NMDAR’s excitotoxicity

Cerebral ischemic injury is characterized by excitotoxicity caused by overactivation of NMDARs which leads to mitochondria dysfunction (Schinder et al., 1996), and Ca^2+^ overload results from both excitotoxic and non-excitotoxic Ca^2+^ signaling pathways (Choi, 2020). The excitotoxic glutamate-dependent NMDAR channels have been targets for stroke intervention for over 50 years (Choi, 2020). TRPM2 is one of the glutamate-independent, non-excitotoxic Ca^2+^ channels which has been implicated as a potential target for ischemic stroke in recent years (Tymianski, 2011). The intriguing result in this study is that, as a non-excitotoxic Ca^2+^-permeable channel, TRPM2 exacerbates NMDAR’s excitotoxicity, implying that intervention of TRPM2 may attenuate both non-excitotoxicity as well as excitotoxicity during ischemic stroke.

How could TRPM2 converge the non-excitotoxic Ca^2+^ signaling and NMDAR-dependent excitotoxic Ca^2+^ signaling pathways to mediate neuronal death during MCAO? We revealed that TRPM2 physically interacts with GluN2a and GluN2b in both heterologous-expressing HEK293T cells and in mouse brains. The physical interaction leads to enhanced surface expression and increased current amplitude of NMDARs. More importantly, we uncovered that the C-termini of GluN2a and GluN2b bind to the EE_3_ domain at the N-terminus of TRPM2, which is a stretch of 16 residues containing three “EE” repeats separated by five residues. When the physical interaction is disrupted by mutations or truncation of the EE_3_ domain, or by the disrupting peptide TAT-EE_3_, interaction of TRPM2 and NMDARs in HEK-293T cells or in neurons is abolished, leading to elimination of both the enhanced surface expression of NMDARs and its increased channel activity (Figure 2-3).

### TRPM2 controls NMDAR surface expression by interacting with PKCγ during ischemic stroke

NMDARs have been shown to be regulated by PKC through modulating intrinsic channel properties and NMDAR trafficking (Carroll and Zukin, 2002; Lan et al., 2001; Xiong et al., 1998). Interestingly, we found that the neuron-specific isoform of PKC, PKCγ, interacts with TRPM2 through the N-terminal domain of TRPM2. More importantly, the interaction of TRPM2 and PKCγ was significantly increased in MCAO brains *in vivo*, and by H_2_O_2_ in the heterologous expressing HEK293T cells *in vitro*, indicating that oxidative stress and ischemic injury conditions promote TRPM2 and PKCγ association (Figure 4). Consistent with our results, oxidative stress-induced interaction of PKCα with a non-functional shorter isoform of TRPM2 (the alternative slice variant, TRPM2-S) was reported in a previous study, albeit full-length TRPM2 was found not to interact with PKCα (Hecquet et al., 2014). As PKC modulates NMDAR trafficking, it is conceivable that increased interaction of PKCγ and TRPM2 under oxidative stress conditions underlies the mechanism of elevated surface expression of NMDARs in MCAO mouse brains. PKC-mediated NMDAR trafficking to the cell surface has been proposed with different mechanisms. Some studies demonstrated that PKC mediates NMDAR surface trafficking through phosphorylation of serine residues (Ser896 and Ser897) in close proximity to the ER-retention motifs of GluN1 (Horak and Wenthold, 2009; Scott et al., 2001; Standley et al., 2000), whereas others showed that PKC-induced increase of NMDAR activity is not mediated by phosphorylation of NMDARs (Zheng et al., 1999). The latter is supported by another study showing that PKC-induced NMDAR trafficking is mediated by triggering auto-phosphorylation of CaMKII, which is associated with NMDARs (Yan et al., 2011). Nonetheless, it seems that phosphorylation function is important for PKC to mediate NMDAR trafficking regardless of whether it directly phosphorylates NMDARs or indirectly phosphorylates their partners. Indeed, whereas PKCγ co-expressed with TRPM2 and NMDARs caused enhancement of NMDARs surface expression, the dominate-negative PKCγ failed to do so (Figure 4). Moreover, PKC activator PMA induced higher NMDARs surface expression, whereas PKC inhibitor staurosporine blocked PKC-induced effects (Figure 4). The effects of PKCγ in the absence of PMA could be attributed to the basal activity under cell culture conditions when cells are surrounded with various growth factors. As both PKCγ and GluN2a/GluN2b interact with the N-terminus of TRPM2 (Figures 3-4), it is conceivable that under oxidative stress conditions, the increased binding of PKCγ to TRPM2 (Figure 5E-F) may bring PKCγ in closer proximity to NMDARs or their interacting protein partners, thereby increasing PKCγ-induced phosphorylation and subsequent surface trafficking of NMDARs.

Another line of evidence supporting the notion of PKCγ-mediated TRPM2-NMDAR complex trafficking to the cell surface is the inhibitory effects produced by endosidine2, an inhibitor of one component of the exocyst complex (Sans et al., 2003), EXO70 (Zhang et al., 2016). NMDARs interact with the exocyst (Sans et al., 2003) for PKC-induced surface delivery via a secretory pathway (Hirschberg et al., 1998). We found that endosidine2 abolished TRPM2-induced enhancement of NMDAR surface expression, and inhibited NMDAR currents in neurons from WT mice (Figure 4Q-R, Figure S6), suggesting that TRPM2-induced NMDAR surface trafficking involves a PKC activation-induced secretory pathway. Furthermore, disruption of the TRPM2-NMDAR interaction largely eliminated increases in surface expression of NMDARs as well as functional NMDAR currents elicited by PMA (Figure 5), strongly supporting that the PKCγ/TRPM2/NMDAR complex is critical for NMDAR surface trafficking. Although further studies are needed to fully understand the exact mechanism by which PKCγ mediates trafficking of the TRPM2/NMDAR complex, we propose the following working model based on our results. Interactions of TRPM2 with NMDARs and PKCγ create a microenvironment where releasing of NMDARs from ER and trafficking of NMDARs to the cell surface are significantly increased under oxidative stress conditions, conferring enhanced excitotoxicity during cerebral ischemic injury.

### Functional coupling of TRPM2 to NMDARs exacerbates extrasynaptic excitotoxicity during ischemic injury

Our data show that functional coupling between TRPM2 and NMDARs appears to only exacerbate extrasynaptic NMDAR excitotoxicity. Using the disrupting peptide TAT-EE_3_, we found that TAT-EE_3_ not only effectively eliminated the increase of NMDAR surface expression and functional increase of NMDAR currents induced by PKC activation (Figures 5-6), but also sufficiently eliminated OGD-induced excitotoxicity *in vitro* (Figure 6), and significantly attenuated MCAO-induced neuron death *in vivo* (Figure 6). Intriguingly, TAT-EE_3_ prevented the decreases in pERK1/2 and pCREB levels in OGD-treated neurons and MCAO brains, a hallmark of pro-survival signaling pathway which can be shut off by extrasynaptic NMDAR activation during ischemic injury (Hardingham and Bading, 2010; Hardingham et al., 2002). Similar to what we demonstrated, pCREB level can be reduced by more than 50% 24 hrs after MCAO (Zhang et al., 2020). The ability of TAT-EE_3_ and TRPM2 knockout to prevent the decreases in pCREB and pERK1/2 levels in MCAO mice strongly indicates that disruption of TRPM2 and NMDAR coupling largely eliminates extrasynaptic excitotoxicity during ischemic brain injury.

It is not surprising that the TRPM2-NMDAR coupling only exacerbates extrasynaptic NMDAR functions (Figure 6), as TRPM2 is absent from the synapse based on the synaptome databases (Bayes et al., 2012). Other studies have also demonstrated a predominantly extrasynaptic distribution of TRPM2 in cultured hippocampal neurons (Olah et al., 2009). Moreover, a previous study demonstrated that NMDAR expression in hippocampal slices from mice not subjected to MCAO was not influenced by TRPM2-KO (Xie et al., 2011), which is consistent with our results showing that NMDAR surface expression level in sham mice is not different between WT and TRPM2-KO (Figure 1W). Thus, it is unlikely that TRPM2-KO can cause abnormal functions of NMDARs under normal physiological conditions. Indeed, TRPM2-KO mice were behaviorally indistinguishable from WT littermates (Yamamoto et al., 2008), which is consistent with our observation that global TRPM2-KO as well as neuron-specific TRPM2-KO mice behave the same as their WT littermates. Furthermore, TRPM2 only enhanced NMDAR function under ischemic injury conditions, which promotes PKCγ to interact with TRPM2 and subsequently NMDAR trafficking to the cell surface. Therefore, disrupting the interaction between TRPM2 and NMDARs to specifically target extrasynaptic NMDAs and to mitigate ischemic stroke will unlikely generate side effects caused by directly antagonizing NMDARs.

### Disrupting peptide TAT-EE3 protects neurons against ischemic injury in vitro and in vivo

The most exciting result in this study is the effectiveness of the disruption of NMDAR-TRPM2 coupling in protecting mice against ischemic brain stroke. Cell-permeable peptides such as TAT-fused peptides have been well characterized and are considered powerful tools for both medical applications and fundamental basic research (Xie et al., 2020). By disrupting the TRPM2-NMDAR interaction, TAT-EE_3_ effectively eliminated the enhanced extrasynaptic toxicity of NMDARs induced by TRPM2. In neurons treated with 10 μM TAT-EE_3_ and in mice administrated with 100 nmol/Kg TAT-EE_3_, we found that TAT-EE_3_ significantly reduced excitotoxicity induced by OGD, prevented the reduction of pCREB and pERK1/2 level in OGD-treated neurons and MCAO mice, and reduced infarct volume as well as neurological deficits in MCAO mice (Figure 6 and Figure S7). These results indicate that disrupting TRPM2 and NMDAR coupling is a promising therapeutic strategy for ischemic stroke.

Although NMDARs interact with various proteins (Petit-Pedrol and Groc, 2021), including recently reported TRPM4 (Yan et al., 2020), there are several unique aspects about the TRPM2-NMDAR interaction. First, TRPM2 is an oxidative stress-activated Ca^2+^-permeable channel; second, as an ion channel interacting with NMDARs, TRPM2 alters NMDAR surface expression by enhancing NMDAR surface trafficking; third, TRPM2 orchestrates the PLCγ to NMDAR interacting complex to enhance NMDAR surface trafficking; fourth, TRPM2 exacerbates extrasynaptic NMDAR function to increase neuronal death. Since TRPM2 is a non-excitotoxic Ca^2+^-permeable channel, the TRPM2-NMDAR interaction-induced exacerbation of NMDAR excitotoxicity makes TRPM2 a unique molecule that converges non-excitotoxic and excitotoxic Ca^2+^ signaling pathways, which may serve as an effective target for ischemic stroke.

### Translational implications

Our data in this study provide a new strategy of targeting excitotoxicity without directly antagonizing NMDARs. Excitotoxicity by glutamate acting on NMDARs has long been established as the dominant conceptual model underlying neuronal cell death associated with ischemic stroke. However, the outcomes from clinical trials of directly antagonizing NMDARs by NMDAR antagonists have been disappointing (Choi, 2020). The adverse effect of inhibiting NMDAR physiological functions is one of many factors contributing to failed NMDAR antagonists in translational applications. In this study, we show that disrupting the physical and functional coupling of NMDARs and TRPM2 can largely eliminate the extrasynaptic toxicity of NMDARs during ischemic injury while preserving the pro-survival synaptic NMDAR function. Thus, targeting TRPM2 may represent an effective strategy for future design and development of treatments for ischemic stroke.

In summary, we found that TRPM2 in neurons plays a key role in mediating the deleterious effects of TRPM2 in ischemic brain stroke. We uncovered that physical and functional coupling of TRPM2 with NMDARs leads to enhanced extrasynaptic NMDAR toxicity under oxidative stress conditions. We revealed that interaction of TRPM2 with PKCγ underlies the mechanism by which TRPM2 mediates enhancement of NMDAR surface expression and functional increase of NMDAR currents. We identified a specific NMDAR-binding domain of TRPM2, and designed a membrane-permeable disrupting peptide TAT-EE_3_. We demonstrated that uncoupling TRPM2 from NMDARs leads to protective effects *in vitro* and *in vivo*. As TRPM2 is a non-excitotoxic Ca^2+^-permeable channel, our results provide a new conceptual strategy that targeting TRPM2 can eliminate ischemic neuronal toxicity mediated by both non-excitotoxic and excitotoxic Ca^2+^ signaling pathways and protect animals against ischemic stroke.

## STAR * METHODS

Detailed methods are provided in the online version of this paper and include the following:

- KEY RESOURCE TABLE
- CONTACT FOR REAGENT AND RESOURCE SHARING
- EXPERIMENTAL MODEL AND SUBJECT DETAILS

- Animal models
- Cortical neuronal cultures
- Cultured cell lines
- METHOD DETAILS

- Knockout of TRPM2 in mice
- Middle cerebral artery occlusion (MCAO, filament method)
- Neurological deficit score evaluation
- Infarct volume assessment by Triphenyl Tetrazolium chloride (TTC) staining
- Subcloning of TRPM2 and NMDARs
- Neuron isolation, culture and treatment
- *In vitro* oxygen-glucose deprivation model
- Cell culture and transfection
- Real-time monitoring of mitochondrial function
- Ratio calcium imaging experiments
- Surface protein isolation
- Protein extraction and immunoblotting
- Co-immunoprecipitation
- Electrophysiology
- Immunofluorescence staining
- QUANTIFICATION AND STATISTICAL ANALYSIS
- DATA AND SOFTWARE AVAILBILITY

## ACKNOWLEDGMENTS

We would like to thank Drs Rajkumar Verma (UCONN Health) and Louise McCullough (UT Health) for constructive discussions about this project. We thank Dr. Rindy Jaffe for helpful comments to the manuscript. We thank Dr. Andrew M. Scharenberg (University of Washington) for kindly providing TRPM2 plasmid.

This work was partially supported by the National Institute of Health (R01-HL143750) and American Heart Association (19TPA34890022) to LY.

## AUTHOR CONTRIBUTIONS

L.Y. conceived and designed the research. P.Z. designed and performed most of the *in vitro* experiments. Z.Y. and J.F. performed most of the *in vivo* experiments. Z.Y., B.S., Y.H., Z.S., A.S.Y., and J.X. conducted some of the *in vitro* experiments. G.W. conducted MCAO surgeries and supervised others doing *in vivo* MCAO surgeries. B.M. and Y. M. generated knockout mice and provided input to discussion of the research. P.Z. and J.F. helped in preparation of the manuscript. L.Y. wrote the manuscript with help from P. Z. All authors commented on the manuscript.

---

**Supplementary Figure 1.**
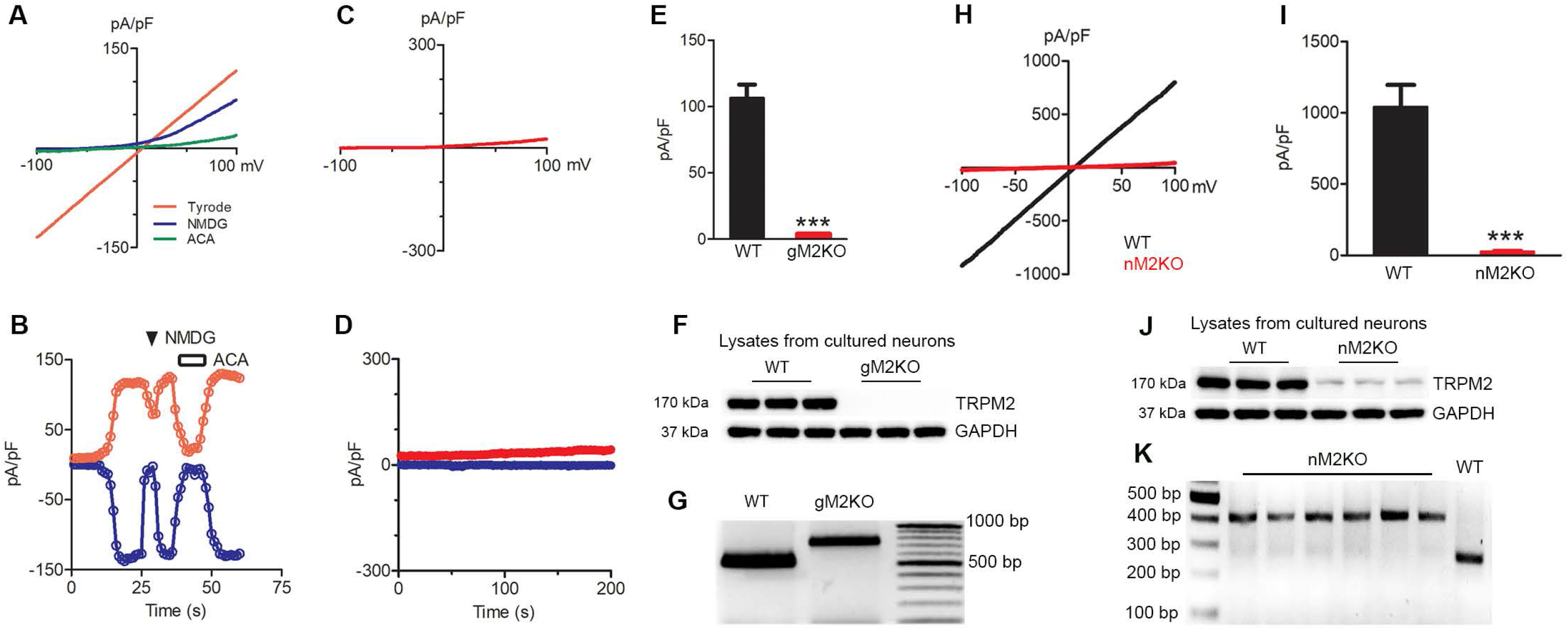
TRPM2 currents recorded in WT neurons and determination of TRPM2 deletion in the global and neuron-specific knockout mice. (A-E), TRPM2 currents recorded in the cortical neurons with pipette solution containing ADPR and Ca^2+^. (A, C), Representative currents elicited by a ramp protocol ranging from −100 to +100 mV in WT neurons (A) but not in global TRPM2-KO (gM2KO) neurons (C). NMDG was used to ensure no leak contamination, and ACA (30 μM) was used to inhibit TRPM2 currents. (B, D), Inward and outward current measured at −100 mV and +100 mV were plotted against time (B). No currents were recorded in the gM2KO neurons (D). (E), Average current amplitude (at +100 mV) of TRPM2 in WT neurons; TRPM2 current was eliminated in gM2KO neurons. Please note that NMDG eliminated inward TRPM2 currents (A, B), indicating no leak current, but meanwhile slightly reduced outward currents (A, B) because elimination of extracellular Ca^2+^ entry will gradually close TRPM2. (F-G), Conformation of global TRPM2 knockout (gM2KO) by genotyping (G) and WB (F). (F), Representative WB results from 3 brains of WT and gM2KO mice. (G), Representative PCR genotyping results showing a 514 bp and 740 bp products for WT and gM2KO mice. (H-I), Representative TRPM2 currents recorded in neurons from WT and neuron-specific TRPM2-KO (nM2KO) mice. TRPM2 currents were recorded in cortical neurons from WT neurons but not in nM2KO neurons (H). Average currents measured at +100 mV (I). Please note that nM2KO eliminated TRPM2 currents. (J-K), Conformation of neuron-specific knockout of TRPM2 by WB and genotyping. (J), Representative WB results to detect TRPM2 deletion using cultured neurons from 3 WT and nestin-cre^+^ floxed mice (nM2KO). TRPM2 protein was largely eliminated in cultured neurons of nM2KO. The trace amounts of protein detected in M2KO neuron cultures is likely from non-neuronal cells in the culture dishes. (K), Representative PCR results for genotyping of TRPM2 flox/flox expression. The predicted PCR products are 400 bp in TRPM2-flox/flox expressing mice (n-6) and 260 bp in a WT mouse as control.

**Supplementary Figure 2.**
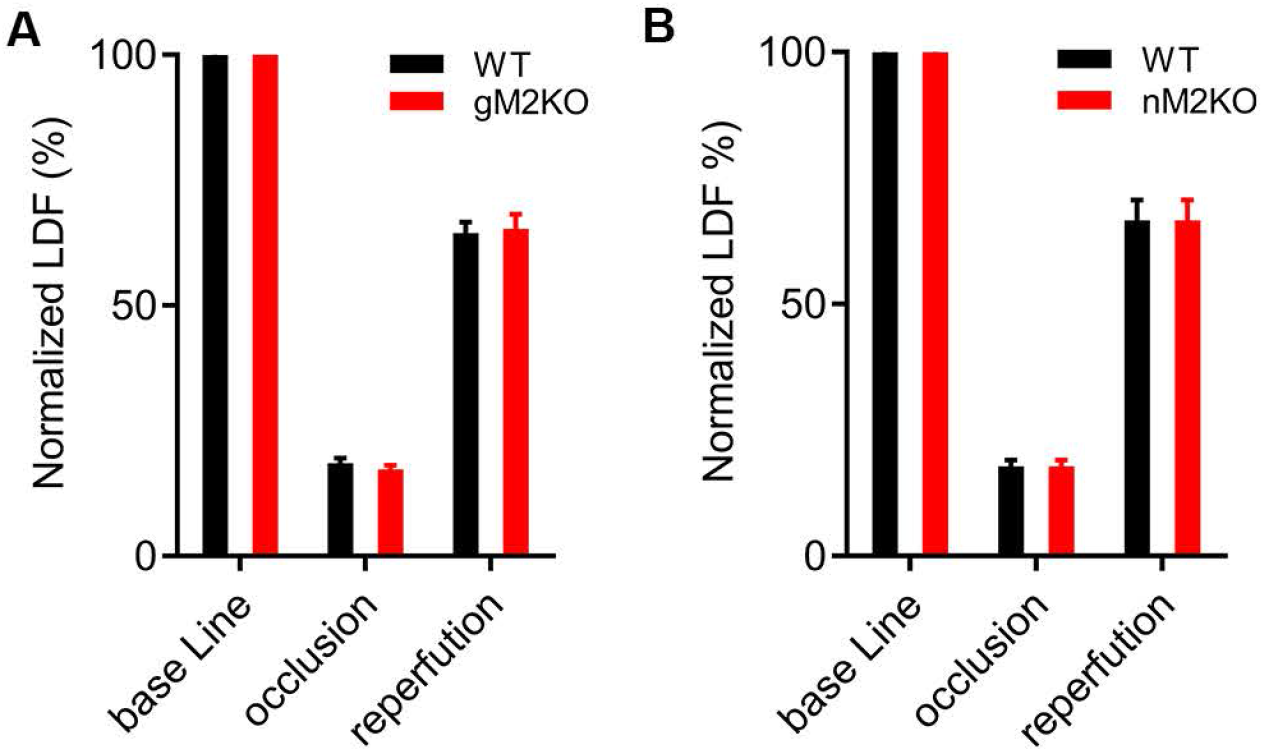
Blood flow changes measured using Laser Doppler Flowmetry (LDF) (A-B), Representative data of blood flow changes measured using LDF. Blood flow was measured using LDF before and after MCAO, as well as after reperfusion. Successful MCA occlusion was confirmed by 85% reduction of cerebral blood flow OGD (***, p < 0.001, unpaired *t*-test, mean ± SEM; n=18 for WT and n=16 for gM2KO groups; and n=13 for WT and n=11 for nM2KO groups).

**Supplementary Figure 3.**
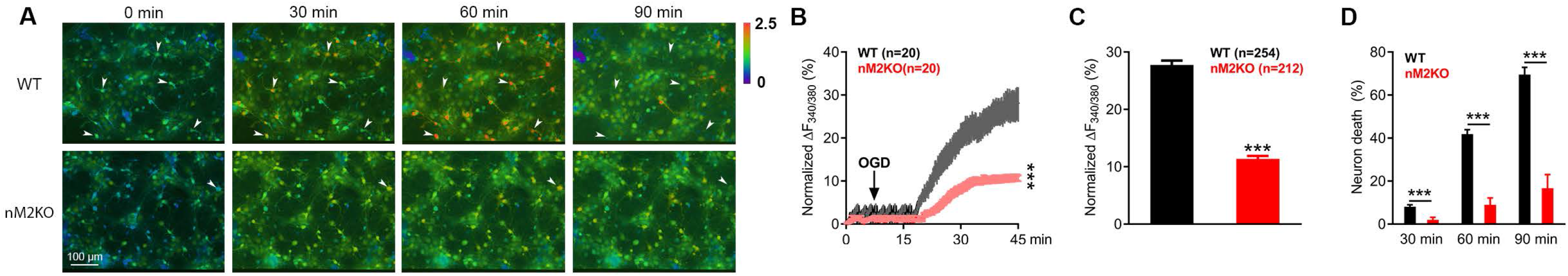
Neuron-specific *Trpm2* Knockout (nM2KO) protects neurons from oxygen-glucose deprivation (OGD)-induced damage. (A), Evaluation of Ca^2+^overload and neuronal death using Fura-2 real-time ratio Ca^2+^ imaging. Cortical neurons were isolated from nM2KO mice (*Trpm2*^flox/flox^, Cre^+^) and WT littermate control mice (*Trpm2*^flox/flox^, Cre^-^) and cultured for 7 to 14 days. Neurons were exposed to OGD and intracellular Ca^2+^ change was monitored by Fura-2 ratio Ca^2+^ imaging for 90 mins. Neurons with increasingly elevated Ca^2+^ levels, such as the ones indicated by arrows, died and disappeared at different time points. Ionomycin was used to induce the maximum Ca^2+^ influx for normalization (not shown). (B), Representative sample traces of Fura-2 real-time Ca^2+^ imaging normalized to ionomycin-induced responses. Averaged traces from 20 neurons which were randomly chosen from WT and nM2KO groups for analysis. Ionomycin was used to induce the maximum Ca^2+^ influx for normalization (***, p < 0.001, unpaired *t*-test, mean ± SEM). (C), Quantification of OGD-induced Ca^2+^ changes after OGD for 30 min. 238 neurons from 3 WT mice in 6 culture dishes and 233 neurons from nM2KO mice in 6 culture dishes were used for analysis (***, p < 0.001, unpaired *t*-test, mean ± SEM). (D), Quantification of OGD-induced neuronal death at 30, 60, and 90 min after OGD (***, p < 0.001, unpaired *t*-test, mean ± SEM). Neuronal death was monitored as F_340/380_ fluorescence gradually reduced and eventually disappeared after the fluorescence reached maximal level (see representative dead cells indicated by arrows in (A)).

**Supplementary Figure 4.**
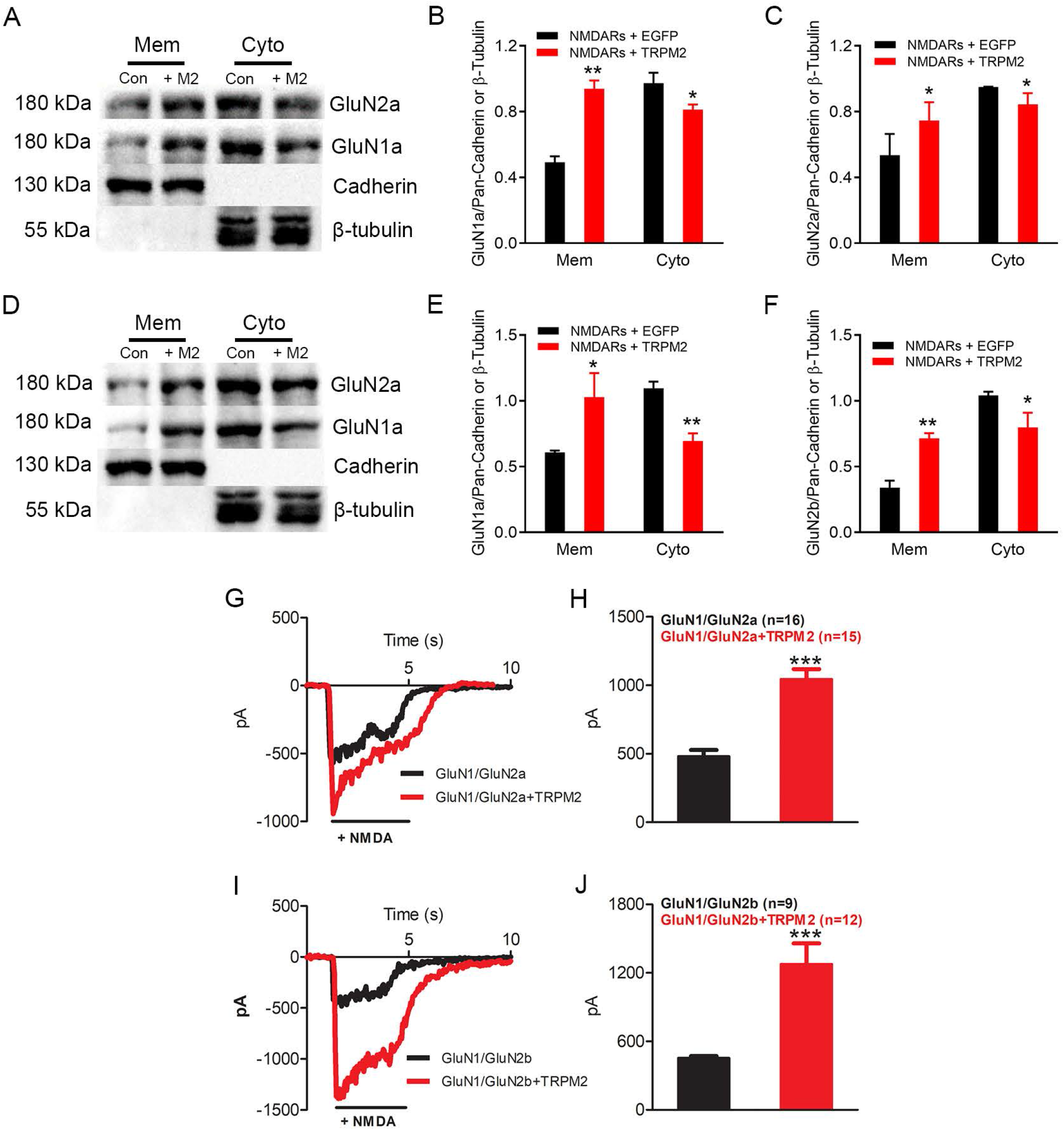
TRPM2 potentiates both GluN1a/GluN2a and GluN1a/GluN2b surface expression levels. (A-C), Surface expression of GluN1a and GluN2a in HEK293T cells co-expressing EGFP (Con) or TRPM2 (+M2). (A), WB analysis of membrane and cytosol levels of GLuN1a and Glu2a. Pan-cadherin and β-tubulin were used as loading controls. (B, C), Quantification of GluN1a, GluN2a membrane (Mem) and cytosol (Cyto) expression (*, p < 0.05, **, p < 0.01; unpaired *t*-test, mean ± SEM, n=3). (D-F) Surface expression of GluN1a and GluN2b in HEK293T cells co-expressing EGFP (Con) or TRPM2 (+M2). (D), WB analysis of membrane and cytosol levels of GluN1a and GluN2b. Pan-cadherin and β-tubulin were used as loading control. (E, F), Quantification of GluN1a, GluN2b membrane (Mem) and cytosol (Cyto) expression (*, p < 0.05, **, p < 0.01; unpaired *t*-test, mean ± SEM, n=3). (G, H) Representative GluN1a/GluN2a currents (G) recorded in HEK-293T cells co-expressing GluN1a/GluN2a/EGFP or GluN1a/GluN2a/TRPM2, and mean current amplitude (H) (***, p < 0.001; ANOVA, Bonferroni’s test; mean ± SEM, n=11∼12). (I, J) Representative GluN1a/GluN2b currents (I) recorded in HEK-293T cells co-expressing GluN1a/GluN2a/EGFP or GluN1a/GluN2a/TRPM2, and mean current amplitude (J) (***, p < 0.001; ANOVA, Bonferroni’s test; mean ± SEM, n=11∼12).

**Supplementary Figure 5.**
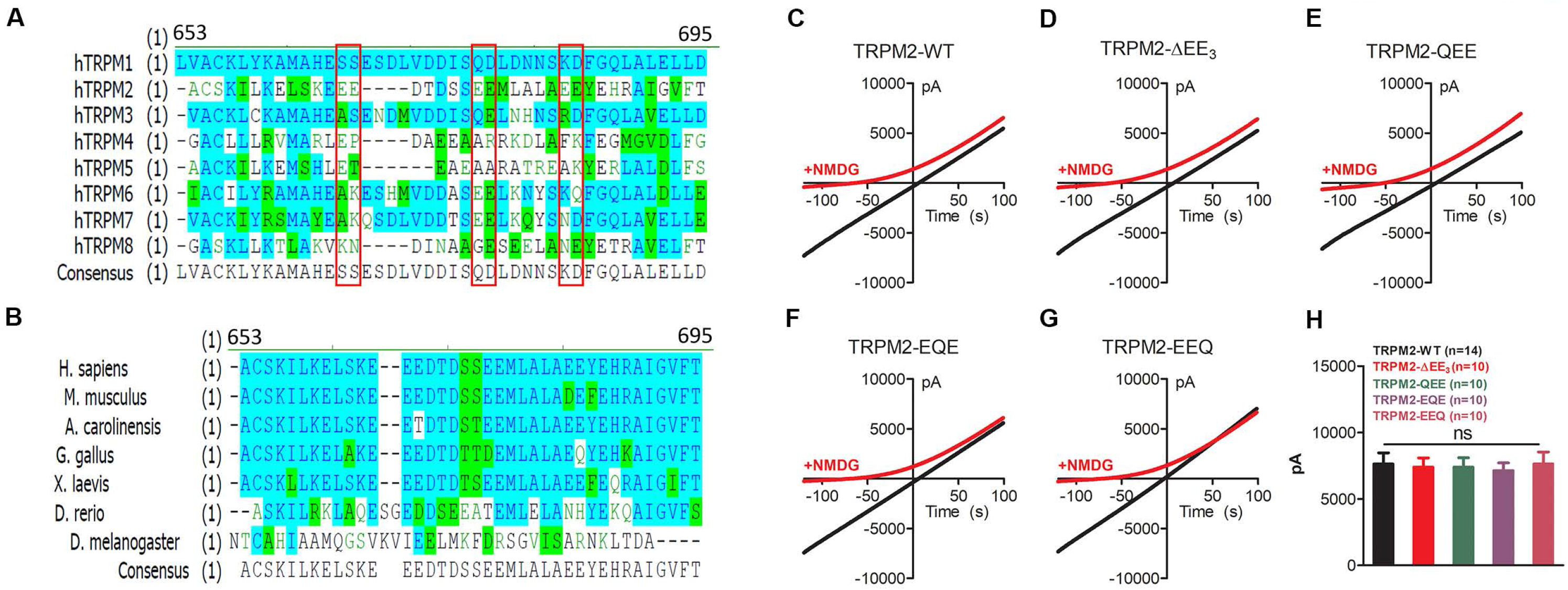
Alignment of triple glutamate-glutamate repeats (EE_3_) and functional evaluation of TRPM2-EE_3_ domain mutants. (A, B) Alignment of triple EE domain (EE_3_) in TRPM subfamily (A), and EE_3_ domain in TRPM2 of different species (B). (C-G) Representative TRPM2 current recordings from HEK-293T cells transfected with EE_3_ domain deleted TRPM2 (TRPM2-ΔEE_3_) and TRPM2 mutants (QEE: E666Q, E667Q; EQE: E673Q, E674Q; EEQ: E680Q, E681Q). (H) Average current quantification of sample traces of TRPM2 current recording from HEK-293T cells transfected with different TRPM2 mutants. 10 recordings from each group were used for analysis (ns, p>0.05; ANOVA, Bonferroni’s test; mean ± SEM).

**Supplementary Figure 6.**
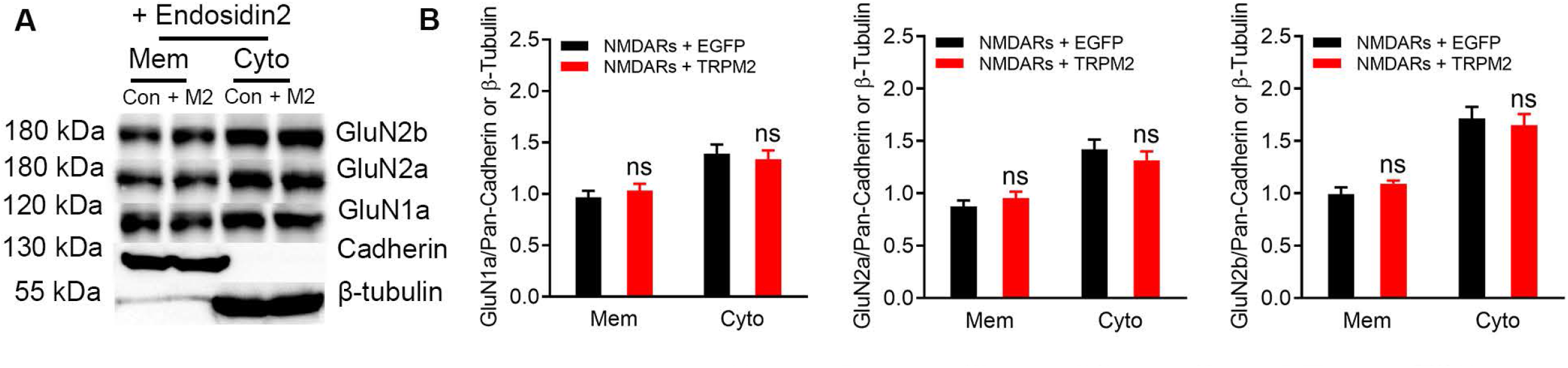
Effects of endosidin2 on surface expression levels of NMDARs. (A), Surface expression of GluN1a, GluN2a, and GluN2b was detected by WB in HEK-293T cells transfected with NMDARs/EGFP (Con) and NMDARs/TRPM2 (+M2) after incubation with 1 µM endosidin2 for overnight. Pan-cadherin and β-tubulin were used as loading control for membrane and cytosolic protein extraction. (B), Quantification of the expression of GluN1a, GluN2a, and GluN2b in cell membrane (Mem) and cytosol (Cyto) (ns, p>0.05; unpaired *t*-test; mean ± SEM, n=3).

**Supplementary Figure 7.**
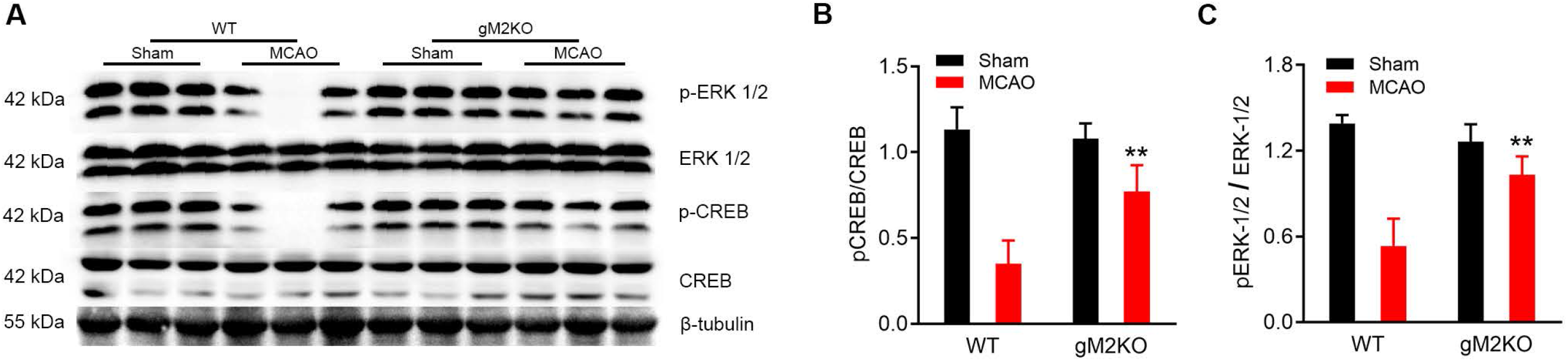
Global TRPM2 Knockout preserves CREB and ERK-1/2 signaling after MCAO. (A), Western blotting analysis of changes of p-ERK 1/2, ERK-1/2, p-CREB, and CREB expression in the brain from WT and TRPM2-KO subjected to sham or MCAO surgery. (B-C), Quantification of pERK-1/2 and pCREB after MCAO or sham surgery. Four mice from each group were used for quantification (**, p < 0.01; ANOVA, Bonferroni’s test; mean ± SEM, n=4/group).

## STAR * METHODS

### Table 1 KEY RESOURCES TABLE

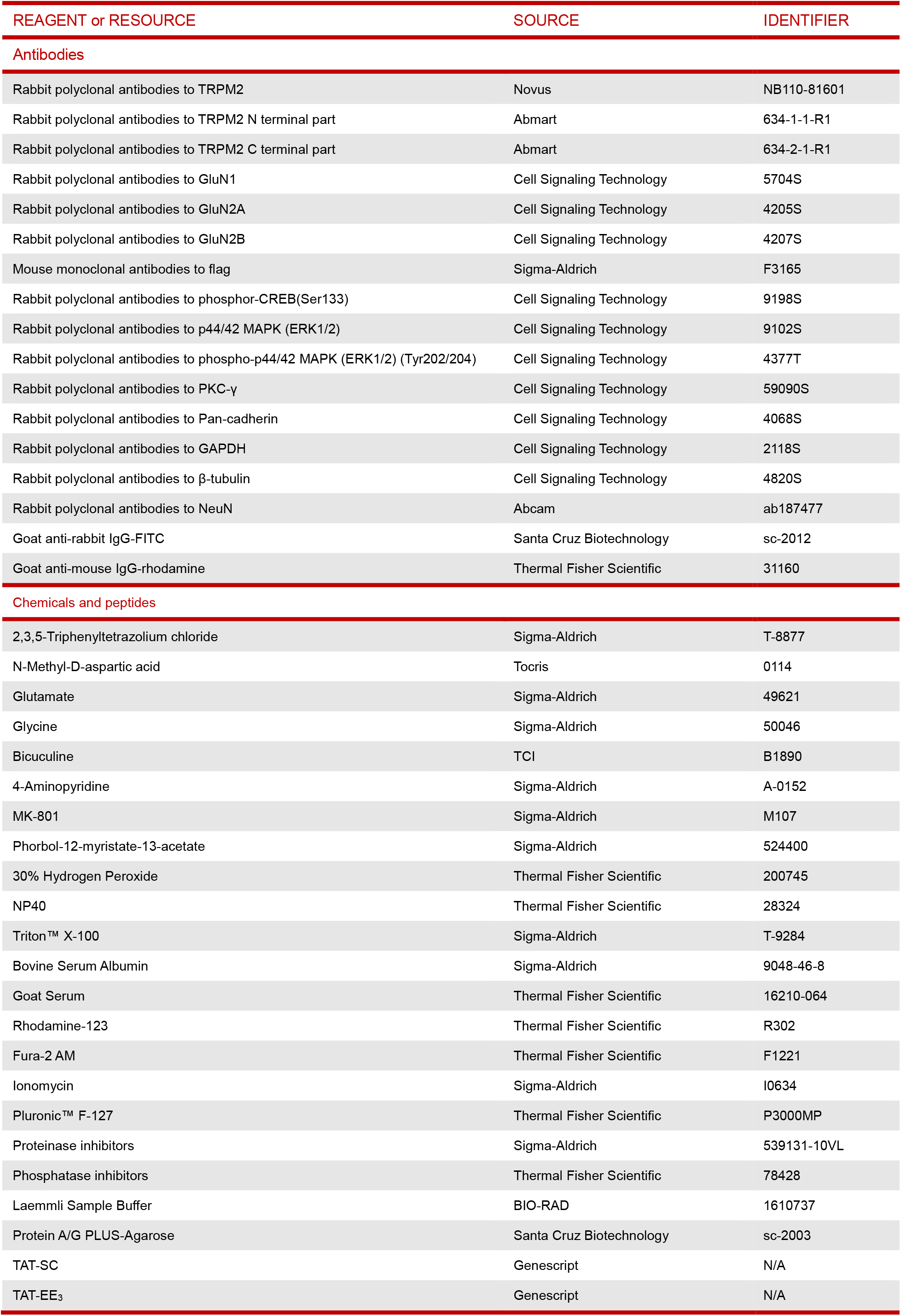
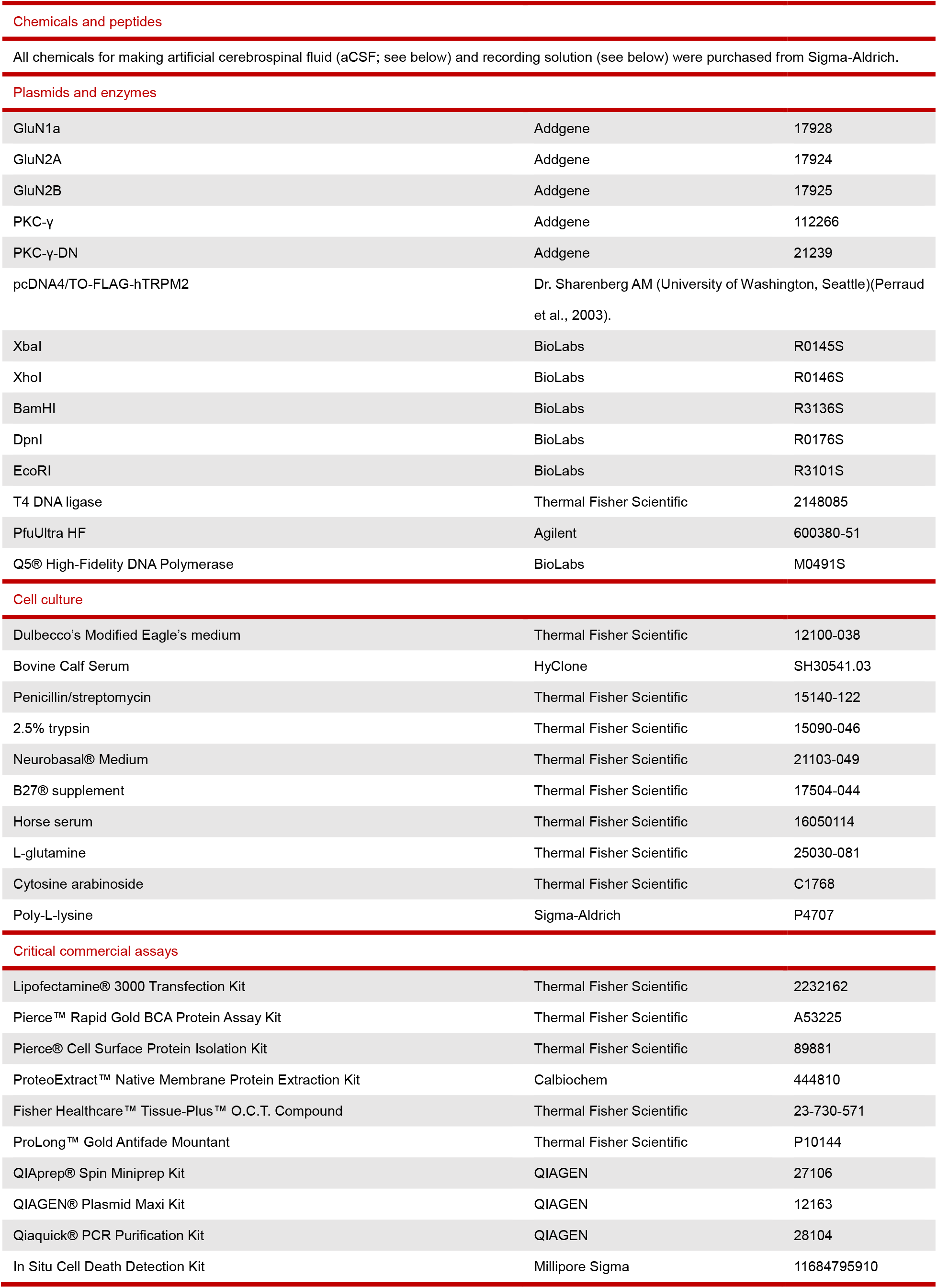
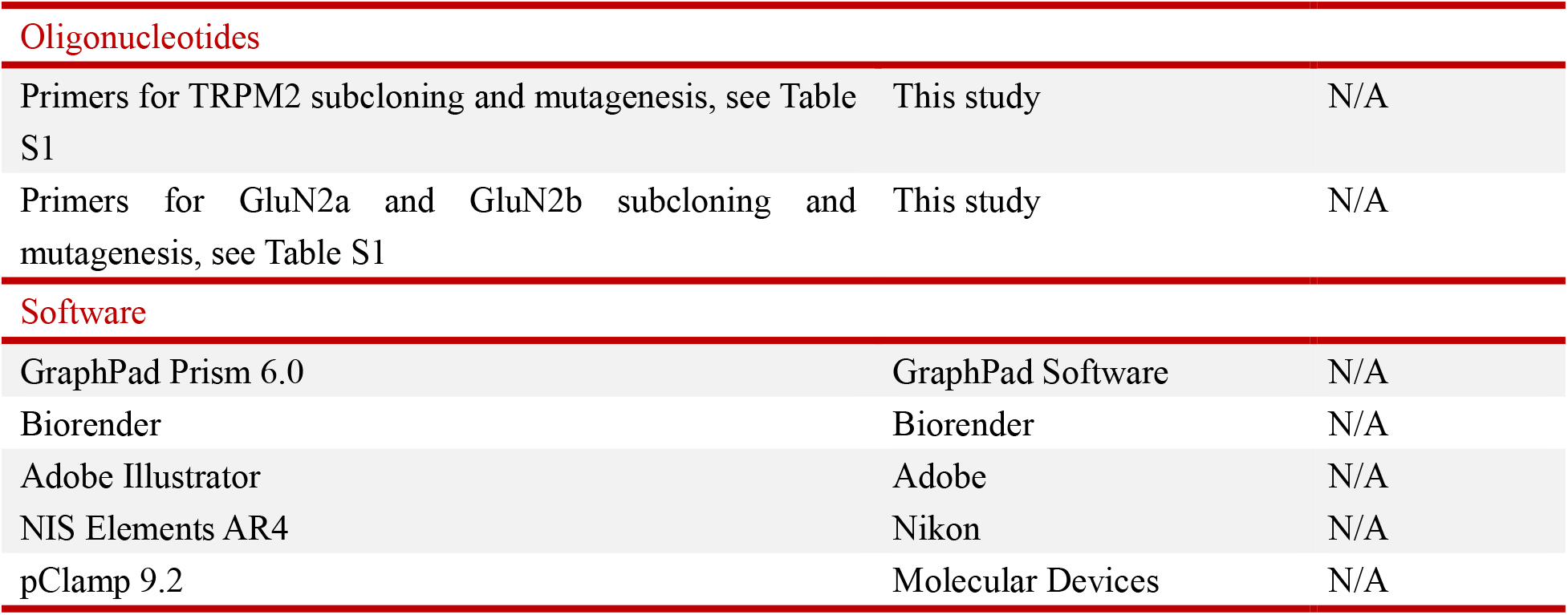

| REAGENT or RESOURCE | SOURCE | IDENTIFIER |
| --- | --- | --- |
| **Antibodies** |  |  |
| Rabbit polyclonal antibodies to TRPM2 | Novus | NB110-81601 |
| Rabbit polyclonal antibodies to TRPM2 N terminal part | Abmart | 634-1-1-R1 |
| Rabbit polyclonal antibodies to TRPM2 C terminal part | Abmart | 634-2-1-R1 |
| Rabbit polyclonal antibodies to GluN1 | Cell Signaling Technology | 5704S |
| Rabbit polyclonal antibodies to GluN2A | Cell Signaling Technology | 4205S |
| Rabbit polyclonal antibodies to GluN2B | Cell Signaling Technology | 4207S |
| Mouse monoclonal antibodies to flag | Sigma-Aldrich | F3165 |
| Rabbit polyclonal antibodies to phosphor-CREB(Ser133) | Cell Signaling Technology | 9198S |
| Rabbit polyclonal antibodies to p44/42 MAPK (ERK1/2) | Cell Signaling Technology | 9102S |
| Rabbit polyclonal antibodies to phospho-p44/42 MAPK (ERK1/2) (Tyr202/204) | Cell Signaling Technology | 4377T |
| Rabbit polyclonal antibodies to PKC-γ | Cell Signaling Technology | 59090S |
| Rabbit polyclonal antibodies to Pan-cadherin | Cell Signaling Technology | 4068S |
| Rabbit polyclonal antibodies to GAPDH | Cell Signaling Technology | 2118S |
| Rabbit polyclonal antibodies to β-tubulin | Cell Signaling Technology | 4820S |
| Rabbit polyclonal antibodies to NeuN | Abcam | ab187477 |
| Goat anti-rabbit IgG-FITC | Santa Cruz Biotechnology | sc-2012 |
| Goat anti-mouse IgG-rhodamine | Thermal Fisher Scientific | 31160 |
| **Chemicals and peptides** |  |  |
| 2,3,5-Triphenyltetrazolium chloride | Sigma-Aldrich | T-8877 |
| N-Methyl-D-aspartic acid | Tocris | 0114 |
| Glutamate | Sigma-Aldrich | 49621 |
| Glycine | Sigma-Aldrich | 50046 |
| Bicuculine | TCI | B1890 |
| 4-Aminopyridine | Sigma-Aldrich | A-0152 |
| MK-801 | Sigma-Aldrich | M107 |
| Phorbol-12-myristate-13-acetate | Sigma-Aldrich | 524400 |
| 30% Hydrogen Peroxide | Thermal Fisher Scientific | 200745 |
| NP40 | Thermal Fisher Scientific | 28324 |
| Triton™ X-100 | Sigma-Aldrich | T-9284 |
| Bovine Serum Albumin | Sigma-Aldrich | 9048-46-8 |
| Goat Serum | Thermal Fisher Scientific | 16210-064 |
| Rhodamine-123 | Thermal Fisher Scientific | R302 |
| Fura-2 AM | Thermal Fisher Scientific | F1221 |
| Ionomycin | Sigma-Aldrich | I0634 |
| Pluronic™ F-127 | Thermal Fisher Scientific | P3000MP |
| Proteinase inhibitors | Sigma-Aldrich | 539131-10VL |
| Phosphatase inhibitors | Thermal Fisher Scientific | 78428 |
| Laemmli Sample Buffer | BIO-RAD | 1610737 |
| Protein A/G PLUS-Agarose | Santa Cruz Biotechnology | sc-2003 |
| TAT-SC | Genescript | N/A |
| TAT-EE<sub>3</sub> | Genescript | N/A |
| **Chemicals and peptides** |  |  |
| All chemicals for making artificial cerebrospinal fluid (aCSF; see below) and recording solution (see below) were purchased from Sigma-Aldrich. |  |  |
| **Plasmids and enzymes** |  |  |
| GluN1a | Addgene | 17928 |
| GluN2A | Addgene | 17924 |
| GluN2B | Addgene | 17925 |
| PKC-γ | Addgene | 112266 |
| PKC-γ-DN | Addgene | 21239 |
| pcDNA4/TO-FLAG-hTRPM2 | Dr. Sharenberg AM (University of Washington, Seattle)(Perraud et al., 2003). |  |
| XbaI | BioLabs | R0145S |
| XhoI | BioLabs | R0146S |
| BamHI | BioLabs | R3136S |
| DpnI | BioLabs | R0176S |
| EcoRI | BioLabs | R3101S |
| T4 DNA ligase | Thermal Fisher Scientific | 2148085 |
| PfuUltra HF | Agilent | 600380-51 |
| Q5® High-Fidelity DNA Polymerase | BioLabs | M0491S |
| **Cell culture** |  |  |
| Dulbecco's Modified Eagle's medium | Thermal Fisher Scientific | 12100-038 |
| Bovine Calf Serum | HyClone | SH30541.03 |
| Penicillin/streptomycin | Thermal Fisher Scientific | 15140-122 |
| 2.5% trypsin | Thermal Fisher Scientific | 15090-046 |
| Neurobasal® Medium | Thermal Fisher Scientific | 21103-049 |
| B27® supplement | Thermal Fisher Scientific | 17504-044 |
| Horse serum | Thermal Fisher Scientific | 16050114 |
| L-glutamine | Thermal Fisher Scientific | 25030-081 |
| Cytosine arabinoside | Thermal Fisher Scientific | C1768 |
| Poly-L-lysine | Sigma-Aldrich | P4707 |
| **Critical commercial assays** |  |  |
| Lipofectamine® 3000 Transfection Kit | Thermal Fisher Scientific | 2232162 |
| Pierce™ Rapid Gold BCA Protein Assay Kit | Thermal Fisher Scientific | A53225 |
| Pierce® Cell Surface Protein Isolation Kit | Thermal Fisher Scientific | 89881 |
| ProteoExtract™ Native Membrane Protein Extraction Kit | Calbiochem | 444810 |
| Fisher Healthcare™ Tissue-Plus™ O.C.T. Compound | Thermal Fisher Scientific | 23-730-571 |
| ProLong™ Gold Antifade Mountant | Thermal Fisher Scientific | P10144 |
| QIAprep® Spin Miniprep Kit | QIAGEN | 27106 |
| QIAGEN® Plasmid Maxi Kit | QIAGEN | 12163 |
| Qiaquick® PCR Purification Kit | QIAGEN | 28104 |
| In Situ Cell Death Detection Kit | Millipore Sigma | 11684795910 |

| *Continued* |  |  |
| --- | --- | --- |
| REAGENT or RESOURCE | SOURCE | IDENTIFIER |
| **Oligonucleotides** |  |  |
| Primers for TRPM2 subcloning and mutagenesis, see Table S1 | This study | N/A |
| Primers for GluN2a and GluN2b subcloning and mutagenesis, see Table S1 | This study | N/A |
| **Software** |  |  |
| GraphPad Prism 6.0 | GraphPad Software | N/A |
| Biorender | Biorender | N/A |
| Adobe Illustrator | Adobe | N/A |
| NIS Elements AR4 | Nikon | N/A |
| pClamp 9.2 | Molecular Devices | N/A |

### CONTACT FOR REAGENT AND RESOURCE SHARING

Further information and resources and reagents should be directed to and will be fulfilled by the lead contact, Dr. Lixia Yue

### EXPERIMENTAL MODEL AND SUBJECT DETAILS

#### Animal Models

##### Animal Care

All the experimental mice bred and hosted in the animal facility building of University of Connecticut School of Medicine (UCONN Health) were fed with standard chow diet and water ad libitum. Standard housing conditions were maintained at a controlled temperature with a 12-h light/dark cycle. All experimental procedures and protocols were approved by the Institutional Animal Care and Use Committee (IACUC) of University of Connecticut School of Medicine (animal protocol: AP-200135-0723), and were conducted in accordance with the U.S. National Institutes of Health Guidelines for the Care and Use of Laboratory Animals.

#### Knockout of TRPM2 (TRPM2-KO)

The global TRPM2 knockout (TRPM2-KO, or gM2KO) mice were generated by Dr. Yasuo Mori’s lab at Kyoto University Japan. The deletion of *Trpm2* was developed in C57B6J mouse by replacing the third exon (S5–S6 linker in the pore domain) with a neomycin coding region. The knockout mice exhibited no differences in behavior or impairment in breeding, compared to wild type (WT) C57BJ6 mice (Yamamoto et al., 2008). TRPM2-KO mice were back-crossed to C57BL/6 mice for ≥10 generations before being used for experiments.

The neuron specific knockout of TRPM2 (nM2KO) was generated by breeding TRPM2^fl/fl^ mice with Nestin-Cre ((B6.Cg-Tg)Nes-cre)1kln/J: 003771; JAX laboratory). TRPM2^fl/fl^ mice were generated by Dr. Barbara Miller (Miller et al., 2013) (Penn State University, Pennsylvania). The exons 21 and 22 encoding transmembrane domain 5 and 6 and pore loop were flanked by loxp recombination sites and will be deleted by Cre recombinase (Miller et al., 2013). The mice were backcrossed with C57BL/6 mice for ≥10 generations before being used for experiments. The TRPM2^flox/flox^ (TRPM2^fl/fl^) with Cre^+^ mice and TRPM2^fl/fl^ with Cre^-^ mice from the same litters were paired for experiments throughout the manuscript.

The inducible global knockout was also generated by using TRPM2^fl/fl^ mice breeding with global Cre, Rosa26-CreERT2 (B6.129-Gt(ROSA)26Sor^tm1(cre/ERT2)Tyj^/J: 008463; JAX laboratory). Knockout was induced by Tamoxifen treatment and confirmed by genotyping. The mice were backcrossed with C57BL/6 mice for ≥10 generations before being used for experiments.

#### Middle cerebral artery occlusion (MCAO)

Eight- to nine-week-old male mice (∼25g) were subjected to right middle cerebral artery occlusion (MCAO) for 120 min followed by 24 hours of reperfusion. The genotype information was blinded to the surgeon who conduct the surgeries. MCAO surgery was performed as previously described (Liu and McCullough, 2014; Wu et al., 2012). In brief, mice were anesthetized with 2% isoflurane (vol/vol) in 100% oxygen and the anesthesia was maintained with 1.5% isoflurane during surgery through nose cone (Harvard Apparatus). The unilateral right middle cerebral artery (MCA) occlusion was carried out by advancing a silicone-coated 6-0 monofilament (Doccol Corporation, Sharon, MA) 10 to 11 mm from internal carotid artery bifurcation via an external carotid artery incision (Chiang et al., 2011). Mouse body temperature was monitored by a rectal temperature probe and maintained at ∼ 37°C with an automatic temperature-regulating heating pad connected to animal temperature controller (TCA T-2DF, Physitemp). Cerebral blood flow was monitored after occlusion as well as after reperfusion. The bregma was exposed and the skull bone countersunk at two 3 × 3-mm areas over both MCA supply territories for bilateral monitoring of local cortical blood flow. Successful occlusion was confirmed by 85% reduction of cerebral blood flow monitored by Laser Doppler Flowmetry (LDF) with laser Doppler blood FlowMeter (Moor-VMS-LDF1, Moor Instrument, Dever, UK). Sham control mice underwent the same procedure but without insertion of filament to occlude the MCA.

#### Neurological deficit score evaluation

After 24 hours of MCAO or sham procedures, neurological deficit was scored based on previously reported criteria (Longa et al., 1989). In brief, score 0 is the minimum (best) score representing no neurological deficit; score 1 represents failure to extend left paw; Score 2 represents circling to the left; score 3 represents falling to the left; score 4 represents inability of spontaneously walking and decreased level of consciousness; and score 5 represents death due to brain ischemia. The observer to score the neurological deficit was an experienced observer and blinded by the group assignment and genotype information. If the animal score was 0 or 5, it was removed from the study.

#### Infarct volume assessment by Triphenyl Tetrazolium chloride (TTC) staining

Tetrazolium chloride (Sigma-Aldrich, T-8877) was dissolved in PBS at a concentration of 20 mg/ml 30 min prior to use. Post-stroke mice were euthanized and brains were frozen at −80°C for 5 min, cut into coronary slices at a thickness of 1 mm. Brain slices were stained with 2% TTC (vol/vol) for 20 min, and then washed using PBS for 3 times, and fixed in *10*% *Neutral* buffered *formalin* for later scanning. TTC labels non-injured tissue, leaving the infarct area white. The stained slices were scanned for data analysis using ImageJ software. The infarct volume was calculated and presented as a percentage of total brain volume (Ren et al., 2003).

#### Antibodies, chemicals and reagents

Rabbit polyclonal antibodies to TRPM2 (Novus, NB110-81601, 1:500 in 5% BSA for WB, 1:50 in protein extraction for IP); Rabbit polyclonal antibodies to GluN1 (Cell Signaling Technology, 5704S, 1:1000 in 5% BSA for WB, 1:50 in protein extraction for IP); Rabbit polyclonal antibodies to GluN2A (Cell Signaling Technology, 4205S, 1:1000 in 5% BSA for WB, 1:50 in protein extraction for IP). Rabbit polyclonal antibodies to GluN2B (Cell Signaling Technology, 4207S, 1:1000 in 5% BSA for WB, 1:50 in protein extraction for IP); Mouse polyclonal antibodies to flag (Sigma-Aldrich, F3165, 1:5000 in 5% BSA for WB): Rabbit polyclonal antibodies to GFP (Cell Signaling Technology, 2956S, 1:2000 in 5% BSA for WB); Rabbit polyclonal antibodies to CREB (Cell Signaling Technology, 4820S, 1:2000 in 5% BSA for WB); Rabbit polyclonal antibodies to phosphor-CREB(Ser133) (Cell Signaling Technology, 9198S, 1:2000 in 5% BSA for WB); Rabbit polyclonal antibodies to p44/42 MAPK (ERK1/2) (Cell Signaling Technology, 9102S, 1:2000 in 5% BSA for WB); Rabbit polyclonal antibodies to phospho-p44/42 MAPK (ERK1/2) (Tyr202/204) (Cell Signaling Technology, 4377T, 1:2000 in 5% BSA for WB); Rabbit polyclonal antibodies to PKC-γ (Cell Signaling Technology, 59090S, 1:5000 in 5% BSA for WB); Rabbit polyclonal antibodies to Pan-cadherin (Cell Signaling Technology, 4068S, 1:5000 in 5% BSA for WB); Rabbit polyclonal antibodies to GAPDH (Cell Signaling Technology, 2118S, 1:5000 in 5% BSA for WB); Rabbit polyclonal antibodies to β-tubulin (Cell Signaling Technology, 4820S, 1:5000 in 5% BSA for WB); Mouse monoclonal antibodies to Caspase-3 (Santa Cruz Biotechnology, sc-7272, 1:1000 in 5% BSA for WB, 1:50 in 5% BSA and 15% goat serum for immunofluorescence staining); Rabbit polyclonal antibodies to NeuN (Abcam, ab187477, 1:50 in 5% BSA and 15% goat serum for immunofluorescence staining); Goat anti-rabbit IgG-FITC (Santa Cruz Biotechnology, sc-2012, 1:1000 in 5% BSA and 15% goat serum for immunofluorescence staining); Goat anti-mouse IgG-rhodamine (Thermal Fisher Scientific, 31660, 1:1000 in 5% BSA and 15% goat serum for immunofluorescence staining); Prolong® Gold antifade reagent with DAPI (Life technologies, P36935), HRP-linked anti-rabbit IgG (Cell Signaling Technology, 7074S, 1:10000 in 5% BSA for WB); HRP-linked anti-mouse-IgG (Cell Signaling Technology, 7076S, 1:10000 in 5% BSA for WB); Tetrazolium chloride (Sigma-Aldrich, T-8877); NMDA (Tocris, 0114); Glutamate (Sigma-Aldrich, 49621); Glycine(Sigma-Aldrich, 50046); Bicuculine (TCI, B1890); 4-AP (Sigma-Aldrich, A-0152); MK-801 (Sigma-Aldrich, M107); PMA (Sigma-Aldrich, 524400); 4-α-PMA (Sigma-Aldrich, P128); H_2_O_2_ (Thermal Fisher Scientific, 200745); NP40 (Thermal Fisher Scientific, 28324), Triton™ X-100 (T-9284), Bovine Serum Albumin (Sigma-Aldrich, 9048-46-8), Goat Serum (Thermal Fisher Scientific, 16210-064). All chemicals for making artificial cerebrospinal fluid (aCSF; see below) and recording solution (see below) were purchased from Sigma-Aldrich.

#### Membrane permeable peptide TAT-EE3 for disrupting TRPM2 and NMDARs coupling and scramble control TAT-SC peptides

TAT-SC (sequence: YGRKKRRQRRR VILLKDHTLEYPVF) and TAT-EE_3_ (sequence: YGRKKRRQRRR EEDTDSSEEMLALAEE) were ordered from GenScript Biotech, and dissolved in PBS to make a stock concentration at 10 mM. HEK-293T cells or isolated neurons were treated with TAT-SC or TAT-EE_3_ at a concentration of 10 µM for at least 8h prior to use. Mice were injected with TAT-SC or TAT-EE_3_ intraperitoneally (ip) at a dose of 100 nmol/kg.

#### Plasmids and enzymes

GluN1a (Addgene, 17928), GluN2A (Addgene, 17924), GluN2B (Addgene, 17925), PKC-γ (Addgene, 112266), PKC-γ-DN (Addgene, 21239). The pcDNA4/TO-FLAG-hTRPM2 construct was a kind gift from Dr. Sharenberg AM (University of Washington, Seattle)(Perraud et al., 2003).

XbaI (BioLabs, R0145S), BamHI (BioLabs, R3136S), XhoI (BioLabs, R0146S), DpnI (BioLabs, R0176S), EcoRI (BioLabs, R3101S) and T4 DNA ligase (Thermal Fisher Scientific, 2148085), PfuUltra HF (Agilent, 600380-51), and Q5® High-Fidelity DNA Polymerase (Biolabs, M0491S) were used to generating different deletion or mutation constructs.

#### Subcloning

For TRPM2, subcloning of N terminus (1-727) was achieved by introducing a stop codon (A2282T) by PCR using PfuUltra HF. C terminal of TRPM2 was amplified by PCR using Q5® High-Fidelity DNA Polymerase, cut by EcoRI and XbaI, and inserted into EGFP-C3 vector. To look for the binding part in N terminus of TRPM2, a series of stop codons were introduced by PCR using PfuUltra HF (C1831A, C1994T, G2138T, A2090T). EE_3_ motif was deleted by PCR using Q5® High-Fidelity DNA Polymerase. EE was mutated to QQ by PCR using PfuUltra HF (G2093C, G2096C; G2117C, G2120C; G2138C, G2141C). For GluN2A and GluN2B, C terminal was amplified by PCR using Q5® High-Fidelity DNA Polymerase, cut by EcoRI and XbaI, and inserted into EGFP-C3 vector. Deletion of C terminus of GluN2A and GluN2B was achieved by introducing a stop codon by PCR using PfuUltra HF (G4371T for GluN2B and G4518T for GluN2B). The information of all the primers is listed in the Table S1.

#### Cell culture and transfection

HEK293T cells were cultured in Dulbecco’s Modified Eagle’s medium (DMEM) (Thermal Fisher Scientific, 12100-038) supplemented with 10% BGS (HyClone, SH30541.03) and 0.5% penicillin/streptomycin (Thermal Fisher Scientific, 15140-122) at 37 °C and 5% CO2. 8h prior to transfection, culture medium was replaced with DMEM supplemented only with 2.5% BGS. Cells were transfected when at a confluence about 80-90% using Lipofectamine® 3000 Transfection Kit (Thermal Fisher Scientific, 2232162) based on instruction.

#### Neuron isolation and culture

Mice pups at P0 were euthanized based on animal protocol. Whole brain was dissected out immediately and immersed in ice-cold Hank’s Balanced Salt Solution (HBSS). Meninges were removed thoroughly, and tissue of different brain areas was taken based on purposes. Brain tissue was cut into small pieces and digested with 0.25% trypsin (Thermal Fisher Scientific, 15090-046) in HBSS at 37 °C for 20 min. Digestion solution was quickly removed, and tissue pellets are washed with Neurobasal® Medium (Thermal Fisher Scientific, 21103-049) for 3 times. Cells were resuspended with appropriate amount of Neurobasal® Medium supplemented with 2% B27® supplement (Thermal Fisher Scientific, 17504-044), 3% horse serum (Thermal Fisher Scientific, 16050114), 0.25% L-glutamine (Thermal Fisher Scientific, 25030-081) and 1% penicillin/streptomycin (Thermal Fisher Scientific, 15140-122). Isolated cells were counted and plated on coverslips pre-coated with poly-L-lysine (Sigma-Aldrich, P4707) at a density about 500 x 10^3^ cells/cm^2^ for OGD and H_2_O_2_ treatment, and 100 x 10^3^ cells/cm^2^ for current recording. Cytosine arabinoside (Sigma-Aldrich, C1768) was added to maintain a concentration at 1µM to inhibit the proliferation of non-neuronal cells. 24 h after plating, culture medium was changed to Neurobasal® Medium supplemented with 2% B27® supplement, 0.25% L-glutamine and 1% penicillin/streptomycin. The concentration of Cytosine arabinoside (araC) was increased to 2 µM. Medium was changed every 3 days. OGD and H_2_O_2_ treatment was conducted at 7^th^ day of culture, and current recording was conducted at 7^th^, 10^th^ and 14^th^ day of culture.

#### Oxygen-glucose deprivation

Oxygen-glucose deprivation (OGD) was achieved by replace the glucose in aCSF with sucrose, and 95% N_2_ and 5% CO_2_ was used to equilibrate sucrose-aCSF to displace oxygen. This condition typically yielded a pO2 of < 5 mm Hg in the imaging chamber (Thompson et al., 2006). At least 10 min was allowed for neurons to adapt to the change from culture medium to aCSF before OGD was applied.

#### Real-time monitoring of mitochondrial function

Mitochondria function was evaluated using Rhodamine-123 dequenching as previously reported. Rhodamine-123 (Rh123,Thermal Fisher Scientific, R302) was dissolved in DMSO to make a stock concentration at 10 mg/ml. Pre-warmed Neurobasal® Medium was used to dilute Rhodamine-123 to 5 µg/ml as working concentration. Culture medium was removed and cultured neurons on the 25 mm coverslip were washed using prewarmed PBS for 3 times, then 2 ml of Rh123 working solution was added. Cells were incubated with Rh123 at 37 °C for 5 min. Then Rh123 working solution was replaced with culture medium. At least 10 min were allowed to achieve Rh123 equilibration after the transition of culture medium to aCSF before experiments.

Fluorescence intensities at 509 nm with excitation at 488nm was collected every 15 s for 30 min using CoolSNAP HQ2 (Photometrics) and data were analyzed using NIS-Elements (Nikon).

#### Ratio calcium imaging experiments

Changes of intracellular Ca^2+^ was measured using ratio Ca^2+^ imaging as we described previously (Du et al., 2010). In brief, Fura-2 AM (Thermal Fisher Scientific, F1221) was dissolved in DMSO to make a stock concentration at 1 mM. Pre-warmed Neurobasal® Medium (Thermal Fisher Scientific, 21103-049) was used to dilute Fura-2 AM to a working concentration at 2.5 µM, and 0.02% Pluronic™ F-127 (Thermal Fisher Scientific, P3000MP) was added to facilitate loading of Fura-2 AM. Cells plated on 25 mm glass coverslips were washed using pre-warmed PBS for 3 times, and then incubated with 2 ml of Fura-2 AM working solution for 30∼45 min at 37 °C. Non-incorporated dye was washed away using HEPES-buffered Saline Solution (HBSS) containing (in mM): 20 HEPES, 10 glucose, 1.2 MgCl_2_, 1.2 KH_2_PO_4_, 4.7 KCl, 140 NaCl, 1.3 Ca^2+^ (pH 7.4).

Ca^2+^ influx was measured by perfusing the cells with Tyrode’s solution for transfected HEK293T cells or aCSF for neurons under different conditions. Ionomycin (Iono) at 1 μM was applied at the end of the experiment as an internal control. Fluorescence intensities at 510 nm with 340 nm and 380 nm excitation were collected at a rate of 1 Hz using CoolSNAP HQ2 (Photometrics) and data were analyzed using NIS-Elements (Nikon). The 340:380 nm ratio in the presence of different treatments was normalized to the maximal Ca^2+^ signal elicited by 1 μM Ionomycin (Iono) as we previously reported (Du et al., 2010).

#### Co-immunoprecipitation

NP-40 lysis buffer (10% NP40, 150 mM NaCl, 1 mM EDTA, 50 mM Tris, pH=8.0) containing proteinase inhibitors (Sigma-Aldrich, 539131-10VL) and phosphatase inhibitors (Thermal Fisher Scientific, 78428) was used to lyse both cultured cells and frozen brain tissue. For transfected cells, proteins were extracted 36 hours after transfection. Cell and tissue lysate were lysed by ultrasound using an ultrasonic cleaner (Thermal Fisher Scientific) filled with ice-cold water for 30 min. After incubated on ice for 1 h, lysate was centrifuged at 13000 g for 30 min and supernatant was collected. Protein concentration was measured using Pierce™ Rapid Gold BCA Protein Assay Kit (Thermal Fisher Scientific, A53225). 300 µg of protein was taken and diluted using NP-40 lysis buffer to make a total volume of 500 µl. Unused protein was allocated and frozen at −80 °C for future use. Appropriate amount of antibody was added based on instruction. After protein-antibody mixture was incubated on ice for 2 h, 25 µl of pre-washed Protein A/G PLUS-Agarose (Santa Cruz Biotechnology, sc-2003) was added, and the whole mixture was incubated at 4 °C for overnight. Then the mixture was centrifuged at 2500g for 1min to get agarose beads. Agarose beads was washed using NP-40 lysis buffer for 7 times, mixed with same amount of 2x Laemmli Sample Buffer (BIO-RAD, 1610737), and boiled at 95 °C for 5 min. Then samples were ready for western blotting analysis.

#### Western blotting

NP-40/Triton lysis buffer (10% NP40, 1% Triton™ X-100, 150 mM NaCl, 1 mM EDTA, 50 mM Tris, pH=8.0) containing proteinase inhibitors and phosphatase inhibitors was used to lyse both cultured cells and frozen brain tissue. Surface protein was extracted using Pierce® Cell Surface Protein Isolation Kit (Thermal Fisher Scientific, 89881) in transfected HEK-293T cells, and using ProteoExtract™ Native Membrane Protein Extraction Kit (Calbiochem, 444810) in brain tissue based on instructions. For transfected cells, proteins were extracted 36 hours after transfection. Cell and tissue lysate were lysed by ultrasound using an ultrasonic cleaner filled with ice-cold water for 30 min. After incubated on ice for 1 h, lysate was centrifuged at 13000 g for 30 min and supernatant was collected. Protein concentration was measured using Pierce™ Rapid Gold BCA Protein Assay Kit.

30-50 µg of total protein was loaded and separated proteins were transferred to Nitrocellulose membranes. Membranes were blocked with 5% BSA and 2.5% goat serum in Tris buffered saline (TBS, pH=7.4) at room temperature for 2 h, and incubated with primary antibodies in TBS with 0.05% Tween (TBS-T) at room temperature for 2 h. Then membranes were incubated with secondary antibodies in TBS-T for 1 h at room temperature for 1 h for detection. Blots were developed with ImageQuant LAS 4000 imaging system. Band intensity was quantified using ImageJ software and normalized with appropriate loading controls.

#### Electrophysiology

Whole cell currents were recorded using an Axopatch 200B amplifier. Data were digitized at 10 or 20 kHz and digitally filtered offline at 1 kHz. Patch electrodes were pulled from borosilicate glass and fire-polished to a resistance of ∼3 MΩ when filled with internal solutions. Series resistance (R_s_) was compensated up to 90% to reduce series resistance errors to <5 mV. Cells in which R_s_ was >10 MΩ were discarded (Du et al., 2009b).

For heterologous expression, transfected HEK-293 cells were identified by GFP fluorescence. TRPM2 current recording in transfected HEK-293T cells was performed as we previously reported (Du et al., 2009a, b). TRPM2 and NMDAR currents recordings from cultured neurons were performed using sCSF as extracellular solution as we previously reported (Zeng et al., 2010). In brief, for TRPM2 current recordings, voltage stimuli lasting 250 ms were delivered at 1-s intervals, with voltage ramps ranging from −100 to +100 mV at holding potential of 0 mV to elicited currents. For NMDA current recordings, a gap-free protocol at holding potential of −80 mV was applied to elicit NMDA currents upon agonist stimulation. A fast perfusion system was used to exchange extracellular solutions and to deliver agonists and antagonists to the cells, with a complete solution exchange achieved in about 1–3 s (Jiang et al., 2005).

Normal Tyrode solution contained (mM): 145 NaCl, 5 KCl, 2 CaCl_2_, 10 HEPES, 10 glucose, osmolarity=290-320 mOsm/Kg, and pH=7.4 was adjusted with NaOH. Extracellular solution for current recording in neuron, the aCSF, solution contained (mM):124 NaCl, 2.5 KCl, 2 MgSO_4_, 2 CaCl_2_, 1.2 NaH_2_PO_4_, 24 NaHCO_3_, 5 HEPES, 12.5 glucose, osmolarity=300-310 mOsm/Kg, with pH=7.4 adjusted with NaOH. For oxygen-glucose-deprivation (OGD) solution, glucose was eliminate from extracellular solution, and the solution was saturated with nitrogen (N_2_) bubbling for 30 min before the experiments.

The internal pipette solution for whole cell current recordings of TRPM2 contained (in mM): 135 Cs-methanesulfonate (CsSO_3_CH_3_), 8 NaCl, 0.5 CaCl_2_, 1 EGTA, and 10 HEPES, with pH adjusted to 7.2 with CsOH. Free [Ca^2+^]_i_ buffered by EGTA was 100 nM calculated using Max chelator (Du et al., 2009b). ADPR 200 μM was included in the pipette solution for most experiments. The intracellular pipette solution to test the effects of OGD on TRPM2 currents in neuron was adjusted to sub-optimal condition, containing (in mM) 135 CsCH_3_SO_4_, 8 NaCl, 0.01 CaCl_2_, 1 MgCl_2_, 10 HEPES (pH 7.2) and 10 μM ADPR.

The intracellular solution for NMDAR current recording contained (mM): 110 K-ASP, 20 KCl, 1 MgSO_4_, 0.05 mM EGTA-K^+^, 0.1 GTP, 5 ATP-Mg_2_, 10 HEPES, osmolarity=275-285 mOsm/Kg, pH=7.2 adjusted with KOH. For the experiments using cells pretreated with the disrupting peptides TAT-SC and TAT-EE_3_, 10 µM TAT-SC or TAT-EE_3_ was included in the pipette solution, and at least 10 min was allowed for achieving intracellular equilibration of TAT-SC or TAT-EE_3_ before current recording.

For current recordings in neurons, tetrodotoxin (0.5 μM) was included in the external solution to block voltage-gated Na^+^ current, and 10 μM nifedipine was used to block voltage-gated Ca^2+^ currents for recording TRPM2 currents.

#### Immunofluorescence staining

Brains harvested from mice were frozen at −80 °C prior to use, and was mounted in Fisher Healthcare™ Tissue-Plus™ O.C.T. Compound (Thermal Fisher Scientific, 23-730-571) prior to cutting. Brains were cut into sagittal slices at a thickness of 6 to 8 μm, mounted to Superfrost® Plus Microscope Slides (Thermal Fisher Scientific, 12-550-15), and frozen at −80 °C for future use. Prior to staining, slides were taken to room temperature for at least 30 min allowing for dehydration. Slices were fixed in 10% formaldehyde for 15 min following washing using PBS for 3 times, and incubated in blocking solution containing 5% BSA, 15% goat serum and 1% Triton X-100 at room temperature for 2 h. Primary antibodies were diluted as described previously in TBS-T containing 15% goat serum. Slices were incubated with primary antibodies for at least 12 h at 4 °C following washing using PBS for 3 times, and incubated with secondary antibodies at room temperature for 2 h. Then slices were washed using PBS for 3 times, and mounted using Prolong® Gold anti-fade reagent with DAPI. Slices were kept at 4 °C before taking pictures. TUNEL staining was performed based on the instruction of kit.

#### Data analysis

All data are expressed as mean ± SEM. For two groups’ comparison, statistical significance was determined using Student’s t-test. For multiple groups’ comparison, statistical significance was determined using one-way or two-way analysis of variance (ANOVA) followed by Bonferroni posttest. P<0.05 was regarded as significant.

---

**Table S1.**
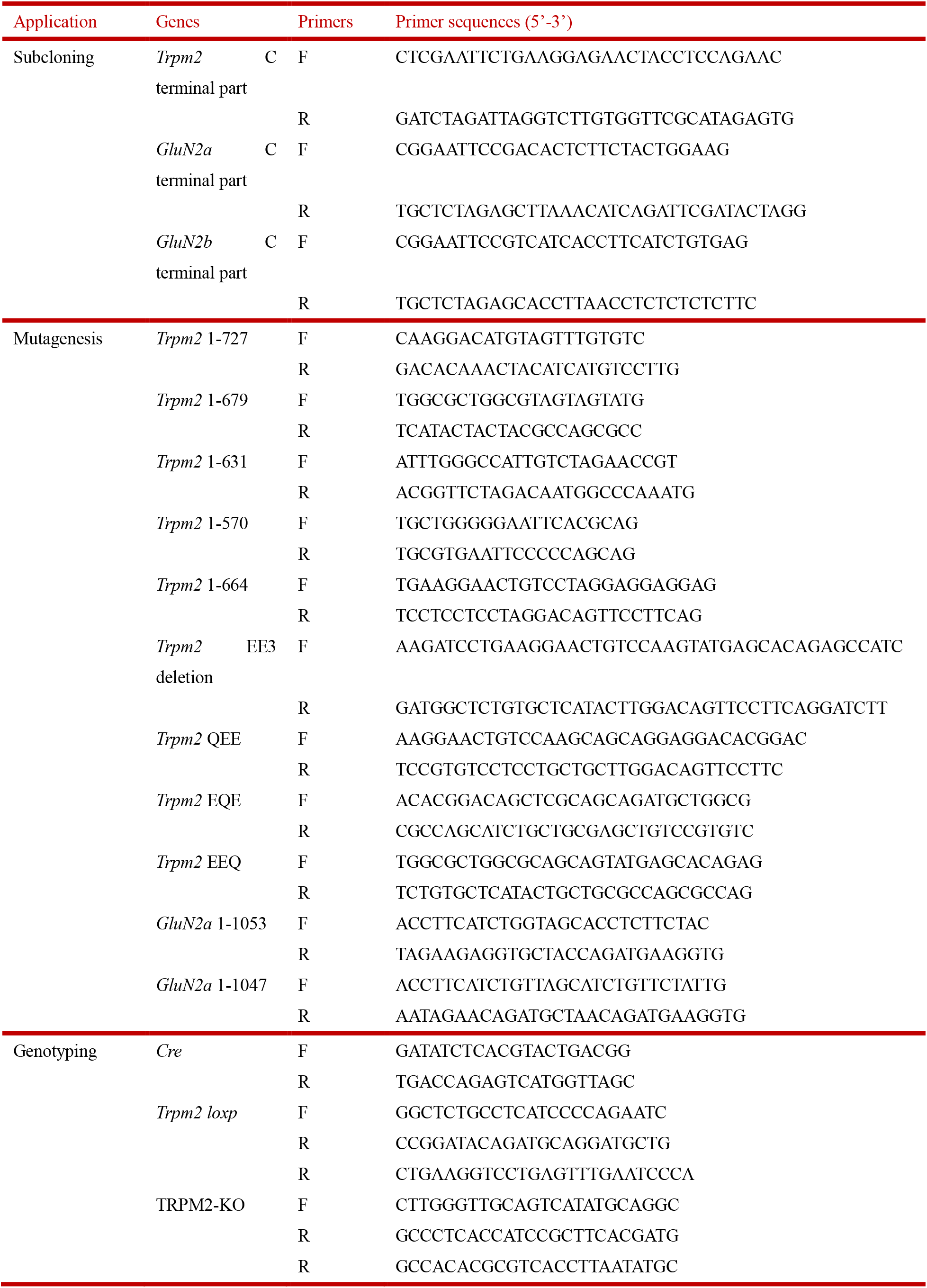
Primers for subcloning, mutagenesis and genotyping.

